# CSF complement proteins are associated with early tau pathology and synaptic damage in an asymptomatic population at risk of Alzheimer’s disease

**DOI:** 10.64898/2026.08.24.742564

**Authors:** Julia Loncke, Mélissa Savard, Cynthia Picard, Henrik Zetterberg, Daniel Auld, Mohamed Badawy, Simon Ducharme, the Alzheimer’s Disease Neuroimaging Initiative, the PREVENT-AD Research Grou, Sylvia Villeneuve, John C. S. Breitner, Judes Poirier

## Abstract

Complement-mediated neuroinflammation has been implicated in Alzheimer’s disease (AD), but its role during the pre-symptomatic phase of the disease remains unclear. In the PREVENT-AD cohort of cognitively unimpaired individuals at increased familial risk of AD, we investigated whether CSF complement proteins relate to early AD pathology and synaptic dysfunction, then assessed our results’ reproducibility across the clinical AD spectrum. Baseline CSF C1q, C3, C3b, and Factor H were measured in relation to CSF AD biomarkers, synaptic proteins, cognition, MRI volumetry, and amyloid and tau PET. Key findings were then examined in 708 participants from ADNI spanning cognitively normal, mild cognitive impairment (MCI), and dementia stages of AD. In PREVENT-AD, C1q was positively associated with CSF P-tau181, T-tau, and multiple synaptic markers including ADAM23, GAP43, SNAP25, and SYT1. Factor H showed similarly strong positive associations with P-tau181, T-tau, ADAM22, ADAM23, GAP43, and SYT1. By contrast, C3 showed minimal associations, while C3b displayed weaker positive relationships with P-tau181, T-tau, ADAM22, and ADAM23. Complement proteins were not robustly associated with amyloid or tau PET, and only C1q related to lower global cognitive performance. In ADNI, C1q emerged as the most consistent analyte, showing positive associations with tau, neurofilament light, and synaptic markers across all diagnostic groups. C3 exhibited predominantly negative associations, whereas C3b and Factor H showed stage-dependent relationships, particularly with evident neurodegeneration and synaptic injury in symptomatic individuals. These findings identify complement dysregulation, especially involving C1q, as an early correlate of tau-linked synaptic pathology, and support a role for complement activation in the AD molecular cascade.

## Introduction

Alzheimer’s disease (AD), the most common form of dementia, manifests as a progressive decline in memory and overall cognition beyond that observed during normal aging, often accompanied by neuropsychiatric disturbances and significant functional impairment.^1,2^ Such symptoms stem from neuropathology characterized by extracellular amyloid plaques, intracellular tau neurofibrillary tangles (NFTs), neuroinflammation, and widespread synaptic dysfunction and loss.^3^ While a small subset of atypical, early-onset cases are linked to familial mutations,^4^ the majority of AD cases appear sporadic and late-onset in nature, with disease processes arising years before symptoms emerge in a preclinical phase of up to two decades.^1,5^ Existing therapies such as newly developed anti-amyloid antibody treatments for individuals with mild cognitive impairment (MCI) and mild AD dementia provide only partial slowing of symptom progression.^6–9^

Recent AD research has emphasized the preclinical phase to elucidate novel biomarkers of early disease mechanisms that may guide the development of preventive interventions. Because synaptic dysfunction, degeneration, and loss correlate strongly with cognitive decline in AD,^10–12^ measurable markers of synaptic pathology are of potential interest for this purpose, along with better-known indicators of amyloid and, especially, tau pathology.^13,14^ In this context, we note recent reports that synapses in autopsied-AD brains are packed with tau oligomers, most notably when persons with neurofibrillary tau pathology have substantial cognitive symptoms.^15^ Previous studies have also reported elevated CSF levels of synaptic proteins such as growth-associated protein 43 (GAP43),^16^ synaptosomal-associated protein 25 (SNAP25),^17^ and synaptotagmin-1 (SYT1)^18^ in those with AD dementia compared to age-matched controls. Furthermore, soluble oligomeric amyloid-beta (Aβ) and phosphorylated tau (P-tau181) have been shown to exert direct synaptotoxic effects independent of later fibrillar or aggregated states.^15,19^ Recent work by our group further demonstrated that CSF markers of both pre- and post-synaptic dysfunction were positively associated with early tau pathology and cognitive deficit in an asymptomatic population at familial risk of AD,^20^ providing evidence for ongoing synaptic degeneration early in the preclinical stages of the disease. Synapses therefore represent a primary site of pathological convergence in AD, particularly in relation to tau pathology.

Mounting research has also pointed toward neuroinflammation as a strong correlate of AD pathogenesis,^3,21,22^ in that inflammatory processes intersect directly with synaptic dysfunction. Specifically, the complement system has emerged as a key mediator linking neuroinflammation to synaptic dysfunction and degeneration.^23^ The complement system comprises a group of serum proteins and cell surface receptors that enhance the function of antibodies and phagocytic cells in the innate immune system,^24,25^ playing a fundamental role in host defence, removal of cellular debris, and the inflammatory response.^26–28^ Within the central nervous system, complement also promotes synaptic maintenance and remodelling, wherein controlled activation of complement pathways promotes microglial-mediated pruning of damaged or redundant synapses during development and plasticity.^28–30^ However, aberrant or excessive complement activation may lead to chronic neuroinflammation and pathological synaptic elimination. Additionally, some data suggest that Aβ may also play a role, such that accumulation of Aβ in AD brain tissue may result in pathological pruning and tau phosphorylation that could translate into neurodegeneration.^31,32^ These findings raise the possibility that synaptic vulnerability in AD is likely driven, at least in part, by immune-mediated mechanisms.

Complement system dysregulation has been increasingly implicated in a variety of neuropsychiatric disorders, particularly those characterized by widespread synaptic dysfunction. Complement mechanisms appear to be underactive in autism spectrum disorder (ASD), as indicated by decreased complement protein and mRNA levels in the brain.^33^ Conversely, in schizophrenia, *C4* gene overexpression has been consistently linked to excessive complement activation and synaptic elimination^34,35^, with elevated levels of C4-derived proteins reported in the CSF of patients.^36^ Similar mechanisms of complement overactivation have also been extensively observed in AD, wherein elevated levels of primarily classical (C1q, C3, C4) but also alternative (C3, Factor H) complement pathway proteins have been measured.^37–39^ Moreover, classical––but not alternative––pathway complement proteins have been found to colocalize with both amyloid plaques and neurofibrillary tangles in post-mortem AD brain tissue.^40–43^ Finally, the 75 genome-wide significant AD risk loci identified to date include complement genes *CR1* and *C1S*.^44,45^

In sum, these findings suggest that complement-mediated synaptic degradation by microglia is a convergent mechanism across multiple brain disorders including AD, and requires further characterization. However, investigation of complement dynamics in the preclinical phase of AD, when neuropathological changes are underway, but symptoms are absent, remains lacking. Extending our previous work on the interaction between AD pathology and synaptic proteins, we explored the relationship between complement activation and the clinical and neuropathological features of AD in asymptomatic persons at risk of AD, and in individuals throughout the AD spectrum.

## Materials and Methods

### Pre-symptomatic Study Population: The PREVENT-AD Cohort

#### Demographics

Participants included 386 individuals from the Pre-Symptomatic Evaluation of Experimental or Novel Treatments for Alzheimer’s Disease (PREVENT-AD) cohort at the Douglas Mental Health University Institute in Montréal, Québec (M_age_ = 62.93 ± 5.16 y, 31.29% male, 37.42% *APOE* ε4+) (Table 1). PREVENT□AD is a longitudinal observational cohort of at-risk, cognitively unimpaired (at baseline) adults aged 60 years and older with a parental or multiple-sibling history of AD. Subjects aged 55–59 years were also eligible if they were within 15 years of the age of symptom onset of their youngest affected relative. Upon enrolment, demographic information included, as a minimum, age, sex, *APOE* genotype, and years of formal education. Since 2011, individuals have been followed through annual visits comprising blood and CSF collection, structural and functional neuroimaging, and serial cognitive assessment. Data are archived in PREVENT AD data release 7.0 (https://openpreventad.loris.ca/). Informed consent was obtained from all participants, and all study procedures were approved by the McGill University Faculty of Medicine Institutional Review Board and carried out in compliance with the ethical standards set forth in the Declaration of Helsinki. Detailed recruitment protocol, inclusion and exclusion criteria, and laboratory methods for the PREVENT-AD cohort are published in Tremblay-Mercier et al. (2021)^46^ and Villeneuve et al. (2025).^47^

**Table 1.** Demographics for the PREVENT-AD cohort. *APOE* apolipoprotein E, *CSF* cerebrospinal fluid, *A*β*42* amyloid-beta 42, *P-tau181* phosphorylated tau 181, *T-tau* total tau, *NFL* neurofilament light, *PAR* peak area ratio, *ADAM* a disintegrin and metalloproteinase domain-containing protein, *GAP43* growth-associated protein 43, *NPX* normalized protein expression, *SNAP25* synaptosomal-associated protein of 25 kDa, *SYT1* synatotagmin-1, *MRI* magnetic resonance imaging, *PET* positron emission tomography, *SUVR* standardized uptake value ratio, *ROI* region of interest, *RBANS* Repeatable Battery for the Assessment of Neuropsychological Status. * = *P* < 0.05; ** = *P* < 0.01; *** = *P* < 0.001; **** = *P* < .0001

| N = 163 | Sex |  | Sig. | APOE ε4 Status |  | Sig. |
| --- | --- | --- | --- | --- | --- | --- |
|  | Male<br>(n = 51) | Female<br>(n = 112) |  | ε4–<br>(n = 102) | ε4+<br>(n = 61) |  |
| Age (years) | 62.83 ± 5.20 | 62.98 ± 5.16 |  | 63.69 ± 5.47 | 61.66 ± 4.34 |  |
| CSF Aβ42 (pg/mL) | 1127.87 ± 258.51 | 1204.11 ± 259.13 |  | 1262.89 ± 223.88 | 1029.64 ± 256.95 | **** |
| CSF P-tau181 (pg/mL) | 45.65 ± 12.13 | 45.94 ± 14.95 |  | 834.41 ± 205.12 | 804.19 ± 208.55 |  |
| CSF T-tau (pg/mL) | 258.06 ± 102.35 | 252.54 ± 93.44 |  | 45.66 ± 14.57 | 46.20 ± 13.40 |  |
| CSF NFL (PAR) | 893.13 ± 187.57 | 793.93 ± 207.05 |  | 251.34 ± 94.64 | 259.39 ± 98.97 |  |
| CSF C1q (pg/mL) | 275.10 ± 64.64 | 252.06 ± 64.59 |  | 260.96 ± 66.01 | 255.78 ± 64.45 |  |
| CSF C3 (pg/mL) | 3873.98 ± 1542.37 | 3467.71 ± 1405.97 |  | 3711.30 ± 1489.36 | 3378.06 ± 1381.65 |  |
| CSF C3b (pg/mL) | 405.86 ± 209.35 | 330.81 ± 166.54 |  | 364.80 ± 183.41 | 330.77 ± 180.57 |  |
| CSF Factor H (pg/mL) | 708.76 ± 172.37 | 568.36 ± 157.67 | **** | 608.47 ± 174.32 | 615.52 ± 175.47 |  |
| CSF ADAM22 (NPX) | 8.64 ± 0.12 | 8.62 ± 0.13 |  | 8.64 ± 0.12 | 8.61 ± 0.12 |  |
| CSF ADAM23 (NPX) | 4.24 ± 0.14 | 4.24 ± 0.14 |  | 4.24 ± 0.14 | 4.23 ± 0.15 |  |
| CSF GAP43 (PAR) | 3158.03 ± 966.48 | 2967.87 ± 1037.38 |  | 2994.17 ± 1069.23 | 3105.44 ± 901.69 |  |
| CSF SNAP25 (PAR) | 23.69 ± 6.43 | 21.75 ± 4.99 |  | 21.32 ± 4.97 | 24.52 ± 6.12 |  |
| CSF SYT1 (PAR) | 20.13 ± 3.84 | 19.62 ± 5.14 |  | 19.24 ± 4.72 | 20.97 ± 4.59 |  |
| MRI volume – entorhinal cortex (mm<sup>3</sup>) | 427.49 ± 61.07 | 393.56 ± 56.28 |  | 401.29 ± 58.11 | 409.26 ± 62.71 |  |
| MRI volume – hippocampus (mm<sup>3</sup>) | 6033.90 ± 608.95 | 5700.07 ± 621.47 | ** | 5780.33 ± 645.30 | 5840.87 ± 619.12 |  |
| MRI volume – lateral ventricles (mm<sup>3</sup>) | 25115.51 ± 7864.12 | 21921.58 ± 9034.93 |  | 22725.87 ± 8725.45 | 23124.13 ± 8987.42 |  |
| PET amyloid (SUVR) | 1.16 ± 0.09 | 1.21 ± 0.09 | * | 1.18 ± 0.08 | 1.22 ± 0.11 | * |
| PET tau (SUVR) – metaROI | 1.14 ± 0.06 | 1.14 ± 0.07 |  | 1.14 ± 0.06 | 1.14 ± 0.08 |  |
| PET tau (SUVR) – entorhinal cortex | 1.06 ± 0.10 | 1.05 ± 0.10 |  | 1.06 ± 0.09 | 1.06 ± 0.11 |  |
| PET tau (SUVR) – fusiform gyrus | 1.19 ± 0.07 | 1.20 ± 0.07 |  | 1.20 ± 0.07 | 1.20 ± 0.08 |  |
| PET tau (SUVR) – lingual gyrus | 1.05 ± 0.09 | 1.08 ± 0.07 |  | 1.07 ± 0.08 | 1.07 ± 0.08 |  |
| RBANS – total score | 97.94 ± 8.70 | 102.73 ± 9.02 | ** | 101.26 ± 8.90 | 101.08 ± 9.69 |  |
| RBANS – attention | 104.39 ± 13.74 | 105.63 ± 13.84 |  | 104.97 ± 13.61 | 105.68 ± 14.16 |  |
| RBANS – delayed memory | 100.36 ± 7.23 | 102.64 ± 6.54 |  | 102.42 ± 6.48 | 101.16 ± 7.34 |  |
| RBANS – immediate memory | 98.49 ± 7.52 | 104.68 ± 10.02 | **** | 102.61 ± 9.74 | 102.93 ± 9.76 |  |
| RBANS – language | 97.86 ± 6.39 | 103.39 ± 9.03 | **** | 101.77 ± 8.80 | 101.19 ± 8.40 |  |
| RBANS – visuospatial constructional | 98.37 ± 12.85 | 95.39 ± 12.51 |  | 96.14 ± 11.91 | 96.67 ± 13.93 |  |

#### CSF Analysis

Following an overnight fast, CSF samples were obtained from a subset of 170 PREVENT-AD participants at baseline via lumbar puncture (LP) using a Sprotte 24-gauge atraumatic needle. Samples were centrifuged (≈ 2000 *g*) at room temperature within four hours of collection for 10 minutes to remove cells and insoluble material, then divided into 0.5 mL polypropylene cryotubes and stored at −80 °C. Levels of AD biomarkers Aβ42 and phosphorylated tau 181 (P-tau181), as well as total tau (T-tau), a less specific marker of neurodegeneration, were measured using Innotest enzyme-linked immunosorbent assay (ELISA) kits (Fujirebio, Ghent, Belgium) in accordance with procedures from the Biomarkers for Alzheimer’s Disease and Parkinson’s Disease (BIOMARKAPD) consortium^48^ (Aβ42 Cat. # 81583; P-tau181 Cat. # 81581; T-tau Cat. # 81579). Complement protein levels (C1q, C3, C3b, Factor H) were quantified using MILLIPLEX’s Human Complement Panel 2 (Cat. # HCMP2MAG-19K; MilliporeSigma Canada Ltd., Oakville, ON, Canada) based on Luminex xMAP technology. Synaptic proteins disintegrin and metalloproteinase domain-containing protein 22 (ADAM22) and 23 (ADAM23) were assessed using Olink’s Target 96 Neurology panel, which employs proximity extension assay (PEA) technology (Olink Proteomics AB, Uppsala, Sweden). Synaptic proteins GAP43, SNAP25, and SYT1 were measured using immunoprecipitation followed by liquid chromatography-tandem mass spectrometry (LC-MS/MS) following procedures outlined in Brinkmalm et al. (2014)^17^ and Öhrfelt et al. (2016).^18^ Neurodegenerative marker neurofilament light (NFL) was measured using the Lumipulse immunoassay platform (Fujirebio, Ghent, Belgium).

#### APOE Genotyping

Automated DNA extraction from buffy coat was performed using the QIA Symphony DNA mini kit (Qiagen, Toronto, ON, Canada. *APOE* genotype was determined using the PyroMark Q96 pyrosequencer as described in Quesnel et al. (2024).^49^ Briefly, DNA was amplified using PCR with primers rs429358 amplification forward 5’-ACGGCTGTCCAAGGAGCTG-3’, rs429358 amplification reverse biotinylated 5’-CACCTCGCCGCGGTACTG-3’, rs429358 sequencing 5’-CGGACATGGAGGACG-3’, rs7412 amplification forward 5’-CTCCGCGATGCCGATGAC-3’, rs7412 amplification reverse biotinylated 5’-CCCCGGCCTGGTACACTG-3’ and rs7412 sequencing 5’-CGATGACCTGCAGAAG-3’. *APOE-*ε4 status is categorized as follows: *APOE*-ε4 carriers are defined as having one or two copies of the ε4 allele and labelled as “1”. Non *APOE*-ε4 carriers have no copy of the ε4 allele and are labelled “0”.

#### MRI Brain Volume Extraction

T1-weighted anatomical MRI scans were initially acquired using a 3D MPRAGE sequence on a Siemens TIM Trio 3T scanner using either a 12- or 32-channel head coil, with acquisition parameters TR = 2300 ms, TE = 2.98 ms, TI = 900 ms, flip angle = 9°, 1 mm^3^ isotropic resolution, and a scan time of 5.12 minutes (Siemens Medical Solutions, Erlangen, Germany). Following June 2016, new participants were scanned exclusively with the 32-channel coil to improve signal quality, and the protocol was expanded to include an MP2RAGE sequence (TR = 5000 ms, TE = 2.91 ms, TIs = 700/2500 ms, flip angles = 4°/5°, 1 mm³ resolution, 8.22-minute scan time). This modification enabled quantitative T1 mapping while retaining the original MPRAGE acquisition. Brain cortical volumes were extracted from T1-weighted images using FreeSurfer Version 5.3.0. All native regional brain volumes were adjusted for each participant’s total intracranial volume, standardized using *z* scores. Extreme outlier values (> 3.5 SD) were excluded from analysis. Analysis of volumetric changes were obtained from the subset of participants having both protein and MRI data available (n = 181).

#### PET Imaging Acquisition and Processing

T1-weighted structural MRI scans were first acquired as described above. Duration of scanning sessions varied between 60–90 minutes. Cerebral deposition of Aβ and tau was quantified via PET using flutafuranol (^18^F-NAV4694; Navidea Biopharmaceuticals, Dublin, OH, USA) and flortaucipir (^18^F-AV1451; Eli Lilly Molecular Psychiatry & Company, Indianapolis, IN, USA), respectively. Scans were acquired 40–70 minutes post-injection for amyloid and 80– 100 minutes post-injection for tau. Total amyloid deposition was reported as the average flutafuranol standardized uptake value ratio (SUVR) across all brain regions, while total tau deposition was calculated as the average flortaucipir SUVR across a temporal meta-region of interest (metaROI) encompassing the entorhinal cortex and fusiform, lingual, and parahippocampal gyri.

#### Cognitive Assessment

Cognitive performance was assessed at baseline and each annual visit using the Repeatable Battery for the Assessment of Neuropsychological Status (RBANS). Total and index-specific scores encompassing five cognitive domains of attention, delayed memory, immediate memory, language, and visuospatial constructional ability were recorded. Four equivalent test versions were administered at random in English or French depending on each subject’s preferred language.^50^

### Replication Study Population: The ADNI Cohort

The replication cohort in this study comprised 708 individuals from the Alzheimer’s Disease Neuroimaging Initiative (ADNI), a multi-site, longitudinal, observational study of North American adults aged 55–90 years that combines clinical, biomarker, neuroimaging, and genetic data to examine cognitive and pathophysiological changes throughout the AD spectrum. Participants included 167 cognitively normal (CN) individuals (M_age_ = 74.42 ± 5.94 y, 53.29% male, 24.55% *APOE* ε4+; Table S1), 403 individuals with mild cognitive impairment (MCI) (M_age_ = 72.28 ± 7.55 y, 57.57% male, 52.11% *APOE* ε4+; Table S2), and 138 individuals with diagnosed late-onset sporadic AD (M_age_ = 75.35 ± 8.53 y, 60.87% male, 68.84% *APOE* ε4+, Table S3). At each ADNI study centre, written informed consent was obtained from all research participants. Each study centre received approval from its institutional review board. All research complied with the ethical principles of the Declaration of Helsinki. Data were collected at various participating research sites across North America, including demographic (sex, age, educational attainment), genetic (*APOE* genotype), CSF (proteomics), neuroimaging (MRI, PET), and clinical (cognitive testing) data, and were accessed from the ADNI online database (adni.loni.usc.edu). CSF protein levels were quantified using the SomaScan 7K assay v4.1 (SomaLogic, Inc., Boulder, CO, USA). These CSF biomarkers were not monitored longitudinally; a single time point measurement is available per participant. Detailed methods for the SomaScan 7K measurements have been described previously.^51,52^ Genotyping, neuroimaging, and cognitive assessment methods are also available within the database.

### Statistical Analysis

Welch’s *t* tests were used to compare CSF protein levels, cerebral volumes, amyloid and tau PET, and cognitive test scores as a function of gender and *APOE* ε4 status. Multiple linear regression (MLR) with ordinary least squares (OLS) parameter estimation was then employed to examine the relationship between CSF complement protein levels and i) CSF levels of AD and synaptic biomarkers, ii) cognitive performance, iii) volume of key brain structures implicated in AD, particularly in early disease stages, and iv) cerebral amyloid and tau deposition.

Owing to high positive skewness, certain complement protein levels and brain volume measurements were log-transformed to improve model fits. In structural MRI analyses all volumetric measurements were corrected for intracranial volume (ICV). For PET analyses, both amyloid and tau deposition were quantified via tracer SUVR in the PREVENT-AD cohort. In the ADNI cohort, amyloid deposition was measured in centiloids (CEN) for comparison across amyloid tracers, whereas tau deposition was quantified via SUVR. All regression analyses were adjusted for covariates of sex, age, and *APOE* ε4 carrier status, while cognitive analyses were further adjusted for years of formal education, as well as test version in the case of the RBANS.

All analyses were conducted on baseline data for both cohorts, with the exception of tau PET for the CN and MCI groups in the ADNI cohort, wherein models were also adjusted for year of visit due to insufficient baseline data. Data outliers were identified following the 1.5 × IQR rule and removed prior to analysis. Significance thresholds were set at *P <* .05; in instances where homoscedasticity of model residuals was violated (Breusch-Pagan *P <* .05), HC3 robust standard errors were utilized to compute *P*-values. *P*-values were then corrected for multiple comparisons using the Benjamini-Hochberg (BH) method for false discovery rate (FDR) control. Corrections were applied to families of related tests defined by a common predictor variable, and performed separately within each cohort (PREVENT-AD and ADNI) across corresponding outcome variables. Within ADNI, corrections were further performed separately for each diagnostic group (CN, MCI, and AD). All statistical analyses were performed using R statistical software version 4.3.2 and RStudio version 2024.12.1+563.

## Results

### PREVENT-AD: A Pre-symptomatic Cohort

In the following sections, we report that the most significant finding in the pre-symptomatic PREVENT-AD cohort is that complement C1q tracks strongly with early tau-associated synaptic pathology, more so than with amyloid burden, and well before cognition and imaging show the same degree of consistency. C3, C3b, and Factor H appear to capture different aspects of the tau-synaptic pathology. Total C3 showed little relationship with tau or synaptic markers, whereas C3b showed weaker positive associations with tau and selected synaptic proteins. Factor H, an inhibitory regulator of the alternative complement pathway, was also informative, but in a more complex manner. Factor H was significantly associated with P-tau181 and T-tau, and, to a lesser degree, with some synaptic biomarkers when compared with C1q.

#### Relation of Findings to Demographics

Within the PREVENT-AD cohort, males possessed significantly higher CSF levels of Factor H, as well as larger hippocampal volumes. Females demonstrated significantly greater cerebral amyloid deposition as measured by PET, and better cognitive performance as measured by RBANS immediate memory, language, and total scale scores. Consistent with the literature, *APOE* ε4 carriers possessed significantly lower CSF Aβ42 levels and increased cerebral amyloid deposition (Table 1).

#### Relation to Alzheimer’s Disease Biomarkers

C1q was significantly positively associated with markers of AD neuropathology in the CSF, including Aβ42 (*R*^2^_adj_ = 0.124, *P =* .0008; Fig. 1A), P-tau181 (*R*^2^_adj_ = 0.246, *P <* .0001; Fig. 1B), and T-tau (*R*^2^_adj_ = 0.212, *P <* .0001, Fig. 1C), but not NFL (*P >* .05; Fig. 1D). C3 was not significantly associated with any markers of AD pathology (all *P*s *>* .05; Fig. 1E–H). C3b demonstrated weak but significant positive associations with P-tau181 (*R*^2^_adj_ = 0.076, *P =* .0125; Fig. 1J) and T-tau (*R*^2^_adj_ = 0.106, *P =* .0024; Fig. 1K), but not Aβ42 nor NFL (all *P*s *>* .05; Fig. 1I, L). Factor H exhibited very strong and significant positive associations with P-tau181 (*R*^2^_adj_ = 0.301, *P <* .0001; Fig. 1N) and T-tau (*R*^2^_adj_ = 0.332, *P <* .0001; Fig. 1O). The regression model of Aβ42 versus Factor H reached significance only due to positive main effects of several covariates (Fig. 1M). The association of Factor H with NFL was not significant (*P >* .05; Fig. 1P).

**Figure 1.**
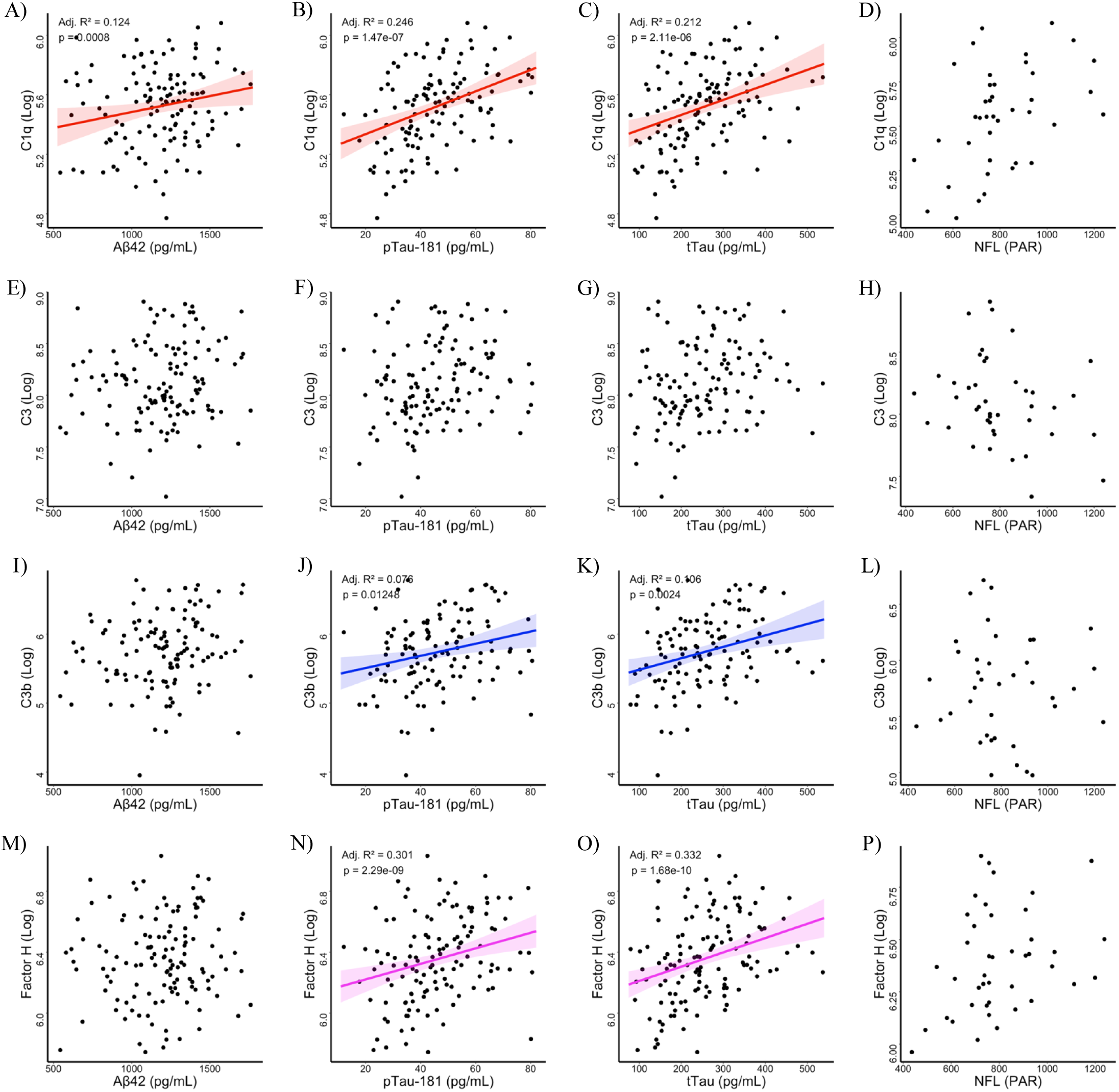
Association between baseline CSF C1q, C3, C3b, and Factor H with AD biomarkers in the PREVENT-AD cohort. CSF complement proteins C1q (**A**, **B**, **C**, **D**; n = 160), C3 (**E**, **F**, **G**, **H**; n = 160), C3b (**I**, **J**, **K**, **L**; n = 158), and Factor H (**M**, **N**, **O**, **P**; n = 160), as well as Aβ42 (**A**, **E**, **I**, **M**; n = 139), P-tau181 (**B**, **F**, **J**, **N**; n = 163), and T-tau (**C**, **G**, **K**, **O**; n = 163), were measured using ELISA. CSF NFL (**D**, **H**, **L**, **P**; n = 49) was measured using immunoprecipitation-LC-MS/MS. Significant linear regressions with a significant main effect of respective AD biomarkers are indicated by a solid fitted line and shaded confidence region. Non-significant linear regression models, as well as significant models owing to main effects of covariates only (age, sex, *APOE* ε4 status), are depicted without fitted lines. Adjusted *R*^2^ and *P*-values of significant models are displayed in the upper left corner of each plot. *R*^2^ values are adjusted for the number of independent variables in each model, while *P*-values are corrected for multiple comparisons using the Benjamini-Hochberg (BH) method for false discovery rate (FDR) control for each dependent variable. PAR = Peak Area Ratio.

#### Synaptic Markers

Again, C1q showed moderate to strong positive associations with pre-synaptic markers of synaptic dysfunction ADAM23 (*R*^2^ = 0.130, *P =* .0005; Fig. 2E), GAP43 (*R*^2^ = 0.307, *P =* .0004; Fig. 2I), SNAP25 (*R*^2^ = 0.222, *P =* .0062; Fig. 2M), and SYT1 (*R*^2^ = 0.284, *P =* .0012; Fig. 2Q). C1q levels only reached significance in relation to postsynaptic marker ADAM22 owing to a positive main effect of age (Fig. 2A). C3 showed a weak positive association with ADAM23 (*R*^2^ = 0.073, *P =* .0109; Fig. 2F), but no other synaptic proteins (all *P*s *>* .05; Fig. 2B, J, N, R). C3b exhibited modest positive associations with ADAM22 (*R*^2^ = 0.110, *P =* .0018; Fig. 2C) and ADAM23 (*R*^2^ = 0.142, *P =* .0004; Fig. 2G), but not with GAP43, SNAP25, nor SYT1 (all *P*s *>* .05; Fig. 2K, O, S). Factor H showed very strong and significant positive associations with ADAM22 (*R*^2^ = 0.255, *P <* .0001 .; Fig. 2D), ADAM23 (*R*^2^ = 0.276, *P <* .0001; Fig. 2H), GAP43 (*R*^2^ = 0.369, *P <* .0001; Fig. 2L), and SYT1 (*R*^2^ = 0.363, *P =* .0001; Fig. 2T). A significant regression model was found for SNAP25 and Factor H, attributed to a positive main effect of male sex (Fig. 2P).

**Figure 2.**
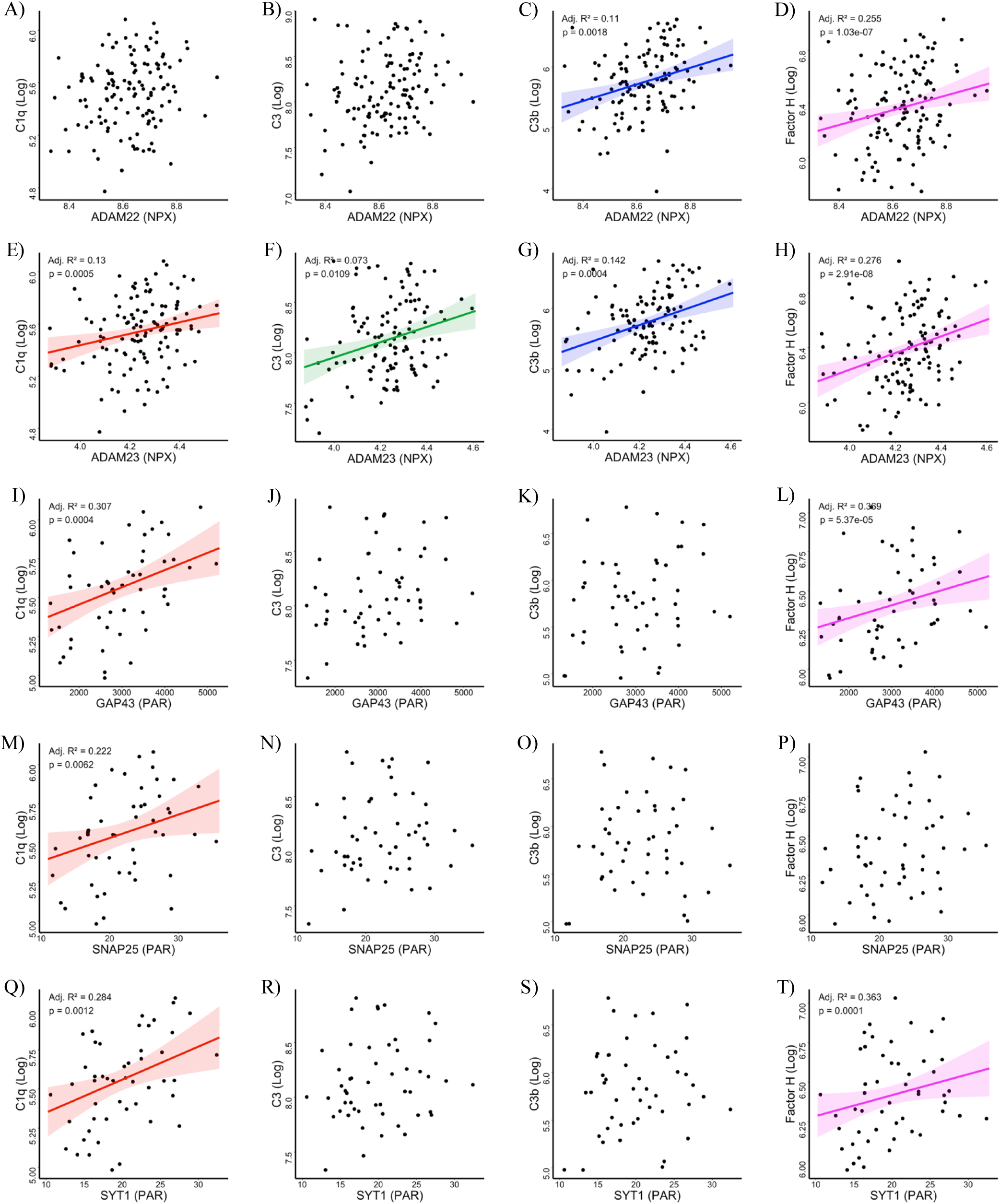
Association between baseline CSF C1q, C3, C3b, and Factor H with synaptic markers in the PREVENT-AD cohort. CSF complement proteins C1q (**A**, **E**, **I**, **M**, **Q**; n = 160), C3 (**B**, **F**, **J**, **N**, **R**; n = 160), C3b (**C**, **G**, **K**, **O**, **S**; n = 158), and Factor H (**D**, **H**, **L**, **P**, **T**; n = 160), were measured using ELISA. ADAM22 (**A**, **B**, **C**, **D**; n = 161) and ADAM23 (**E**, **F**, **G**, **H**; n = 161) were measured using OLINK PEA. GAP43 (**I**, **J**, **K**, **L**; n = 63), SNAP25 (**M**, **N**, **O**, **P**; n = 60), and SYT1 (**Q**, **R**, **S**, **T**; n = 60) were measured using immunoprecipitation-LC-MS/MS. Significant linear regressions with a significant main effect of respective synaptic markers are indicated by a solid fitted line and shaded confidence region. Non-significant linear regression models, as well as significant models owing to main effects of covariates only (age, sex, *APOE* ε4 status), are depicted without fitted lines. Adjusted *R*^2^ and *P*-values of significant models are displayed in the upper left corner of each plot. *R*^2^ values are adjusted for the number of independent variables in each model, while *P*-values are corrected for multiple comparisons using the Benjamini-Hochberg (BH) method for false discovery rate (FDR) control for each dependent variable. NPX = Normalized Protein eXpression; PAR = Peak Area Ratio.

#### Cognitive Performance

RBANS attention index scores correlated negatively with C1q (*R*^2^_adj_ = 0.088, *P =* .0034) and C3b levels (*R*^2^_adj_ = 0.080, *P =* .0082) to a weak degree but only reached significance with C3 and Factor H due to a negative main effect of age. Delayed memory scores exhibited no significant association with any complement protein (all *P*s *>* .05). Similarly, immediate memory scores decreased solely due to negative main effects of covariates but not complement protein levels. Language scores demonstrated a moderate and significant negative association with C1q levels (*R*^2^_adj_ = 0.128, *P =* .0002), but only reached significance with C3, C3b, and Factor H owing to negative main effects of covariates. Visuospatial constructional ability correlated negatively with C3 levels (*R*^2^_adj_ = 0.105, *P =* .0015), but associations with C1q, C3b, and Factor H only reached significance attributable to negative main effects of covariates. Overall, global cognitive performance measured via RBANS total scale score was significantly negatively associated with C1q levels (*R*^2^_adj_ = 0.183, *P <* .0001). In relation to C3, C3b, and Factor H levels, total RBANS scores decreased solely due to negative main effects of covariates (Fig. 3A, Table S4).

**Figure 3.**
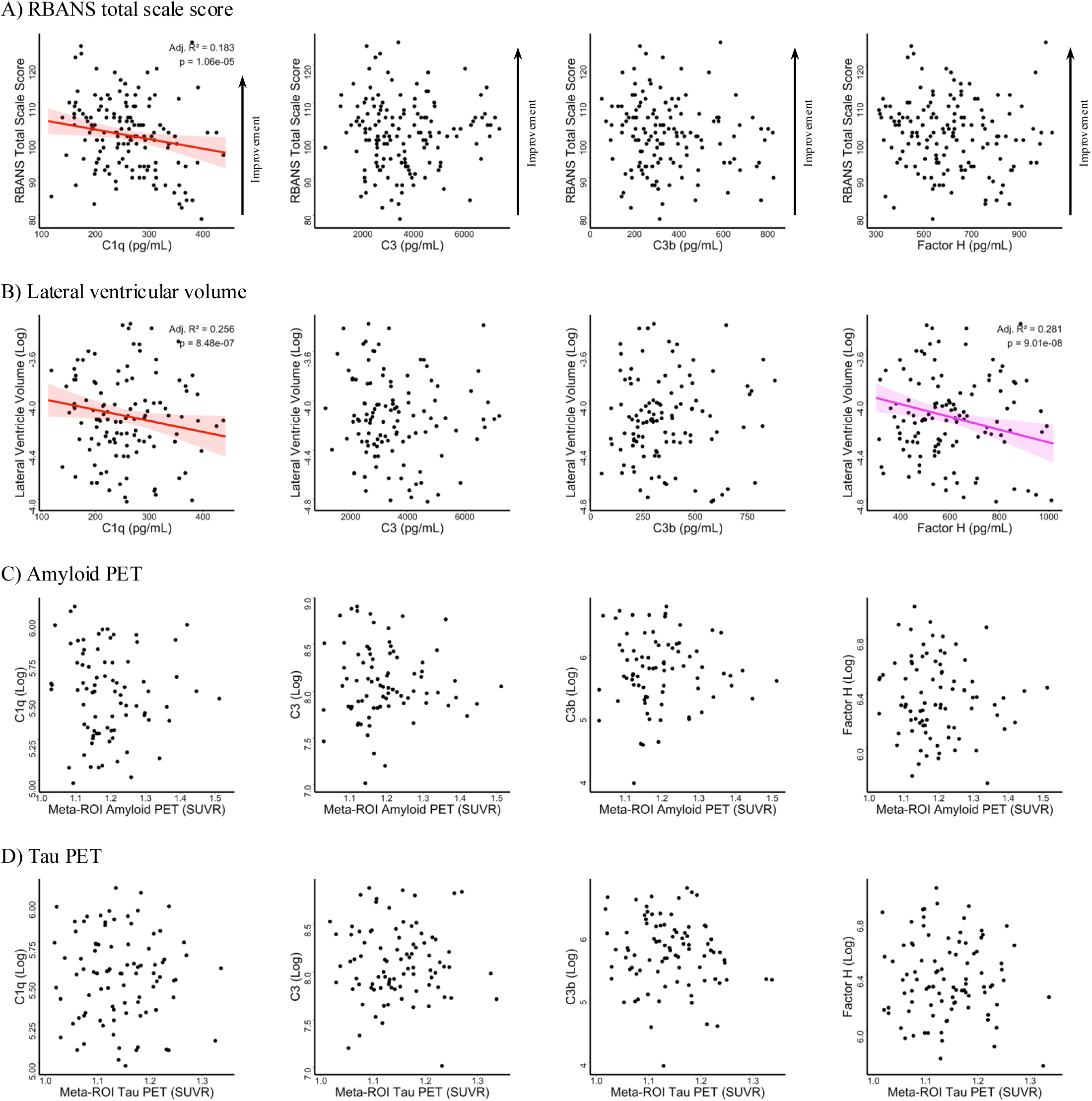
Association between baseline CSF C1q, C3, C3b, and Factor H with cognitive performance, lateral ventricular volume, and amyloid and tau PET at baseline in the PREVENT-AD cohort. CSF complement proteins C1q (n = 160), C3 (n = 160), C3b (n = 158), and Factor H (n = 160) were measured using ELISA (**A**, **B**, **C**, **D**). Complement protein levels were contrasted against cognitive performance, assessed using the Repeatable Battery for the Assessment of Neuropsychological Status (RBANS) (**A,** n = 161) and lateral ventricle volume, measured using volumetric MRI (**B**, n = 156). Amyloid PET (flutafuranol; **C**, n = 119) and tau PET (flortaucipir; **D**, n = 117) were contrasted against complement protein levels. Significant linear regressions with a significant main effect of respective complement proteins or PET are indicated by a solid fitted line and shaded confidence region. Non-significant linear regression models, as well as significant models owing to main effects of covariates only (age, sex, *APOE* ε4 status, with the addition of education for RBANS score), are depicted without fitted lines. Adjusted *R*^2^ and *P*-values of significant models are displayed in the upper right corner of each plot. *R*^2^ values are adjusted for the number of independent variables in each model, while *P*-values are corrected for multiple comparisons using the Benjamini-Hochberg (BH) method for false discovery rate (FDR) control for each dependent variable. SUVR = Standardized Uptake Value Ratio.

#### Cerebral Volumes

No complement protein levels showed any significant association with either left or right entorhinal cortical volume. Regression models for all complement protein levels versus both left and right hippocampal volume reached significance, but these associations were driven by negative main effects of multiple covariates. Curiously, lateral ventricle volume was significantly negatively associated with both C1q (*R*^2^ = 0.256, *P <* .0001) and Factor H levels (*R*^2^ = 0.281, *P <* .0001). Lateral ventricle volume reached significance in regression models with C3 and C3b, again owing to negative main effects of covariates (Fig. 3B, Table S5).

##### Cerebral Amyloid and Tau Deposition

Total cerebral amyloid deposition was not significantly associated with C3 nor C3b levels (all *P*s *>* .05). Associations between these biomarkers and C1q and Factor H levels were due again to positive main effects of covariates (Fig. 3C). The same was true of total tau deposition (Fig. 3D), as well as regional tau deposition in the entorhinal cortex and fusiform and lingual gyri, whereas tau PET showed no significant association with C3 nor C3b, and only reached significance with C1q and Factor H due to positive main effects of covariates (Table S6).

#### ADNI: A Symptomatic Cohort

In ADNI, C1q remained positively associated with P-tau181, T-tau, and synaptic markers across CN, MCI, and AD, replicating the PREVENT-AD observations. In MCI, C1q was the only analyte that showed a convincing positive association with tau PET. In contrast, C3 became more clearly related to NFL in MCI and AD, and to several synaptic markers in later MCI/AD phases. This pattern suggests that activated downstream complement may be more tightly coupled to ongoing neurodegenerative injury once clinical disease is underway. In ADNI, these associations appeared much weaker and shift toward NFL in MCI/AD, with only a modest amyloid PET association in AD.

#### Relation of Findings to Demographics

ADNI participants had an overall mean age of 73.39 ± 7.51 years at baseline, with approximately 57% of participants being male and 49% being *APOE* ε4 carriers across all clinical groups. Among CN participants, males exhibited significantly higher CSF levels of NFL and Factor H, as well as greater hippocampal and lateral ventricular volumes. *APOE* ε4 carriers showed significantly lower CSF levels of Aβ42 and higher CSF levels of P-tau181 (Table S1). Among those with MCI, males demonstrated significantly higher CSF levels of NFL and most complement proteins (C1q, C3b, Factor H), along with greater hippocampal and lateral ventricular volumes. Females exhibited significantly higher CSF levels of C3 and synaptic proteins ADAM23 and SYT1. *APOE* ε4 carriers possessed significantly higher CSF levels of global tau (P-tau181, T-tau), Factor H, and GAP43, but significantly lower levels of NFL and SNAP25. *APOE* ε4 carriers also possessed smaller entorhinal cortex volumes and exhibited increased cerebral amyloid deposition and lower MoCA scores (Table S2). Among participants with AD, males had significantly higher CSF levels of NFL, C3b, and Factor H, whereas females showed higher levels of C3. *APOE* ε4 carriers displayed significantly lower CSF levels of NFL and Factor H, as well as smaller entorhinal cortex volumes (Table S3).

#### Relation to Alzheimer’s Disease Biomarkers

C1q levels demonstrated a moderate and significant positive association with Aβ42 in those with AD (*R*^2^ = 0.149, *P =* .0003) but only reached statistical significance in individuals with MCI due to a positive main effect of age. There was no significant relationship between Aβ42 and C1q levels in CN individuals (*P >* .05). C1q also showed significant positive associations of increasing magnitude with P-tau181 (CN: *R*^2^ = 0.089, *P =* .0026, Fig. 4A; MCI: *R*^2^ = 0.133, *P <* .0001, Fig. 4B; AD: *R*^2^ = 0.157, *P =* .0001, Fig. 4C), T-tau (CN: *R*^2^ = 0.100, *P =* .0011; MCI: *R*^2^ = 0.137, *P <* .0001; AD: *R*^2^ = 0.165, *P <* .0001), and NFL levels (CN: *R*^2^ = 0.106, *P =* .0012; MCI: *R*^2^ = 0.125, *P <* .0001; AD: *R*^2^ = 0.164, *P <* .0001) across clinical groups.

**Figure 4.**
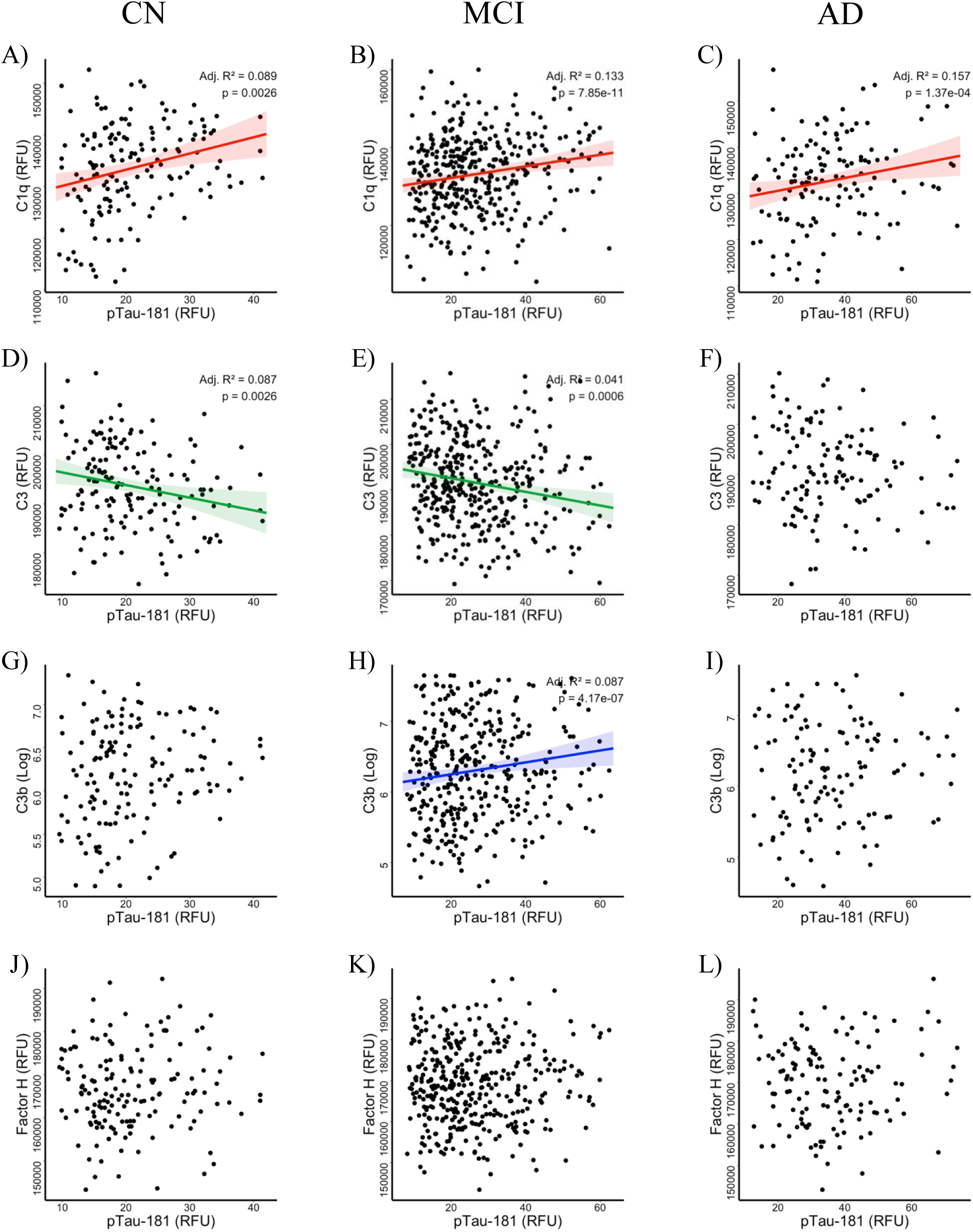
Association between baseline CSF C1q, C3, C3b, and Factor H with P-tau181 in cognitively normal, MCI, and AD individuals from the ADNI cohort. CSF complement proteins C1q (**A**, **B**, **C**; n_CN_ = 167; n_MCI_ = 399, n_AD_ = 138), C3 (**D**, **E**, **F**; n_CN_ = 166; n_MCI_ = 397, n_AD_ = 137), C3b (**G**, **H**, **I**; n_CN_ = 167; n_MCI_ = 402, n_AD_ = 138), and Factor H (**J**, **K**, **L**; n_CN_ = 165; n_MCI_ = 396, n_AD_ = 136), as well as P-tau181 (n_CN_ = 165; n_MCI_ = 402, n_AD_ = 137), were measured using the SomaScan 7K assay. Significant linear regressions with a significant main effect of P-tau181 are indicated by a solid fitted line and shaded confidence region. Non-significant linear regression models, as well as significant models owing to main effects of covariates only (age, sex, *APOE* ε4 status), are depicted without fitted lines. Adjusted *R*^2^ and *P*-values of significant models are displayed in the upper right corner of each plot. *R*^2^ values are adjusted for the number of independent variables in each model, while *P*-values are corrected for multiple comparisons using the Benjamini-Hochberg (BH) method for false discovery rate (FDR) control for each dependent variable within each cognitive diagnostic group. RFU = Relative Fluorescence Units.

Interestingly, C3 levels showed weak but significant negative associations with all markers of AD pathology in CN individuals (Aβ42: *R*^2^_adj_ = 0.114, *P =* .0054; P-tau181: *R*^2^_adj_ = 0.087, *P =* .0026, Fig. 4D; T-tau: *R*^2^_adj_ = 0.098, *P =* .0011; NFL: *R*^2^_adj_ = 0.072, *P =* .0043) and those with MCI (Aβ42: *R*^2^_adj_ = 0.031, *P =* .0056; P-tau181: *R*^2^_adj_ = 0.041, *P =* .0006, Fig. 4E; T-tau: *R*^2^ = 0.051, *P <* .0001; NFL: *R*^2^ = 0.043, *P =* .0004), but only reached significance in AD participants attributable to negative main effects of covariates.

C3b showed no significant association with Aβ42, P-tau181 (Fig. 4G), nor T-tau in CN individuals (all *P*s *>* .05), but displayed a weak positive association with NFL in this group (*R*^2^ = 0.099, *P =* .0042). In those with MCI and AD, the regression models of Aβ42 versus C3b only reached significance due to a main effect of male sex. C3b also exhibited weak positive correlations with P-tau181 (*R*^2^ = 0.087, *P <* .0001; Fig. 4H) and T-tau levels (*R*^2^ = 0.096, *P <* .0001) in MCI subjects, but these models only reached significance in individuals with AD owing to main effects of covariates. However, C3b showed moderate to strong positive associations with NFL levels in MCI (*R*^2^ = 0.119, *P <* .0001) and AD subjects (*R*^2^ = 0.285, *P <* .0001).

Similarly, Factor H was not significantly associated with Aβ42, P-tau181 (Fig. 4J), nor T-tau in CN individuals (all *P*s *>* .05), but displayed a weak positive association with NFL in this group (*R*^2^ = 0.063, *P =* .0066). Regression models of Factor H levels against Aβ42, P-tau181 (Fig. 4K, L), and T-tau only reached significance in the MCI and AD groups due to varying main effects of covariates. Yet, Factor H was significantly positively correlated with NFL levels in MCI (*R*^2^ = 0.057, *P <* .0001) and AD subjects (*R*^2^ = 0.133, *P =* .0003). Full AD biomarker regression statistics for CN (Table S7), MCI (Table S8), and AD (Table S9) participants are provided in the Supplementary Material.

#### Synaptic Markers

CSF C1q levels showed moderate to strong positive associations with synaptic markers ADAM22 (CN: *R*^2^_adj_ = 0.151, *P <* .0001, Fig. 5A; MCI: *R*^2^_adj_ = 0.139, *P <* .0001, Fig. 5B; AD: *R*^2^_adj_ = 0.177, *P <* .0001, Fig. 5C), ADAM23 (CN: *R*^2^_adj_ = 0.122, *P =* .0005, Fig. 5D; MCI: *R*^2^_adj_ = 0.149, *P <* .0001, Fig. 5E; AD: *R*^2^_adj_ = 0.196, *P <* .0001, Fig. 5F), GAP43 (CN: *R*^2^_adj_ = 0.110, *P =* .0023, Fig. 5G; MCI: *R*^2^_adj_ = 0.147, *P <* .0001, Fig. 5H; AD: *R*^2^_adj_ = 0.201, *P <* .0001, Fig. 5I), and SYT1 (CN: *R*^2^_adj_ = 0.116, *P =* .0009, Fig. 5M; MCI: *R*^2^_adj_ = 0.157, *P <* .0001, Fig. 5N; AD: *R*^2^_adj_ = 0.142, *P =* .0004, Fig. 5O) across all clinical groups. C1q levels did not show any association with SNAP25 levels in CN participants (*P >* .05) and only reached significance in MCI and AD individuals due to varying main effects of covariates (Fig. 5J–L).

**Figure 5.**
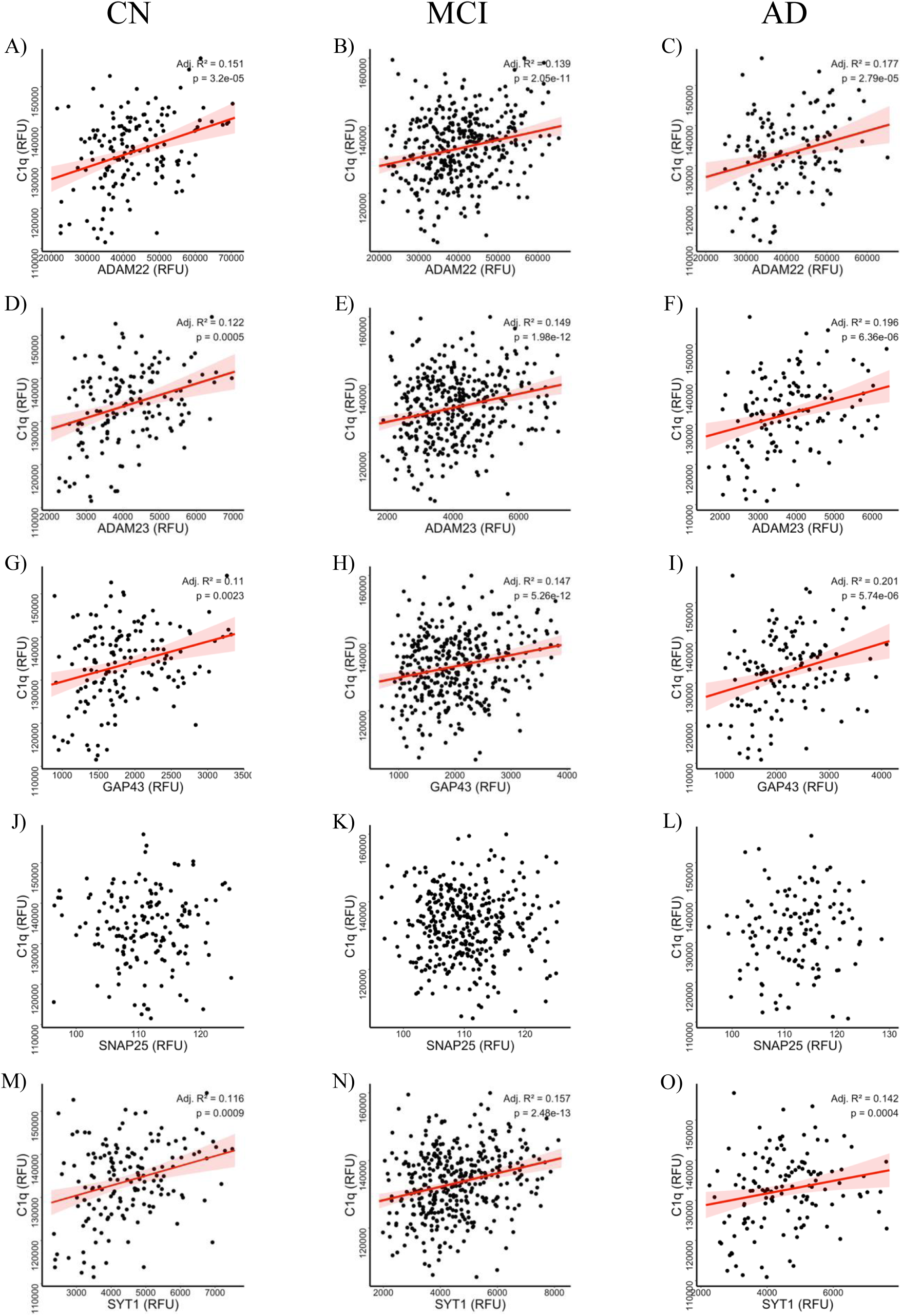
Association between baseline CSF C1q with synaptic markers in cognitively normal, MCI, and AD individuals from the ADNI cohort. CSF complement protein C1q (n_CN_ = 167; n_MCI_ = 399, n_AD_ = 138), as well as synaptic markers ADAM22 (**A**, **B**, **C**; n_CN_ = 165; n_MCI_ = 394, n_AD_ = 135), ADAM23 (**D**, **E**, **F**; n_CN_ = 165; n_MCI_ = 399, n_AD_ = 138), GAP43 (**G**, **H**, **I**; n_CN_ = 166; n_MCI_ = 397, n_AD_ = 138), SNAP25 (**J**, **K**, **L**; n_CN_ =165; n_MCI_ = 389, n_AD_ = 133), and SYT1 (**M**, **N**, **O**; n_CN_ = 166; n_MCI_ = 398, n_AD_ = 137), were measured using the SomaScan 7K assay. Significant linear regressions with a significant main effect of respective synaptic markers are indicated by a solid fitted line and shaded confidence region. Non-significant linear regression models, as well as significant models owing to main effects of covariates only (age, sex, *APOE* ε4 status), are depicted without fitted lines. Adjusted *R*^2^ and *P*-values of significant models are displayed in the upper right corner of each plot. *R*^2^ values are adjusted for the number of independent variables in each model, while *P*-values are corrected for multiple comparisons using the Benjamini-Hochberg (BH) method for false discovery rate (FDR) control for each dependent variable within each cognitive diagnostic group. RFU = Relative Fluorescence Units.

C3 levels showed weak but significant negative associations with most synaptic markers in both CN (ADAM22: *R*^2^ = 0.096, *P =* .0009), (ADAM23: *R*^2^ = 0.082, *P =* .0036), (GAP43: *R*^2^ = 0.087, *P =* .0023), (SYT1: *R*^2^ = 0.096, *P =* .0009) and MCI individuals (ADAM22: *R*^2^ = 0.072, *P <* .0001), (ADAM23: *R*^2^ = 0.080, *P <* .0001), (GAP43: *R*^2^ = 0.053, *P <* .0001), (SYT1: *R*^2^ = 0.071, *P <* .0001). C3 levels did not show any association with SNAP25 in CN participants nor those with MCI (all *P*s *>* .05). In AD subjects, C3 showed weak negative associations with ADAM22 (*R*^2^ = 0.105, *P =* .0011) and SNAP25 levels (*R*^2^ = 0.097, *P =* .0006), but only reached significance in models with ADAM23, GAP43, and SYT1 due to a negative main effect of male sex.

In CN individuals, C3b levels showed no significant correlation with any synaptic markers (all *P*s *>* .05), but displayed weak yet significant positive associations with most synaptic proteins in those with MCI (ADAM22: *R*^2^ = 0.096, *P <* .0001; ADAM23: *R*^2^ = 0.126, *P <* .0001; GAP43: *R*^2^ = 0.093, *P <* .0001; SYT1: *R*^2^ = 0.094, *P <* .0001). The regression model of C3b versus SNAP25 levels in MCI subjects was only significant due to varying main effects of covariates. C3b also showed moderate positive correlations with ADAM22 (*R*^2^ = 0.132, *P =* .0012), SNAP25 (*R*^2^ = 0.169, *P =* .0001), and SYT1 (*R*^2^ = 0.113, *P =* .0029) in those with AD, but only reached significance with ADAM23 and GAP43 due to a positive main effect of male sex.

Factor H levels were not significantly associated with any synaptic protein in CN subjects (all *P*s *>* .05). In those with AD, regression models of Factor H versus synaptic marker levels were only significant due to varying main effects of covariates. In those with MCI, Factor H levels were weakly positively associated with ADAM22 (*R*^2^_adj_ = 0.058, *P <* .0001) and ADAM23 levels (*R*^2^_adj_ = 0.075, *P <* .0001), but were again only significant against GAP43, SNAP25, and SYT1 due to various main effects of covariates. Full synaptic marker regression statistics for CN (Table S10), MCI (Table S11), and AD (Table S12) participants are provided in the Supplementary Material.

#### Cognitive Performance

Across all three clinical groups (CN, MCI (Fig. 6A), AD), regression models relating global cognitive performance via MoCA total scale score to CSF complement protein levels were significant only due to covariate main effects (Table S13).

**Figure 6.**
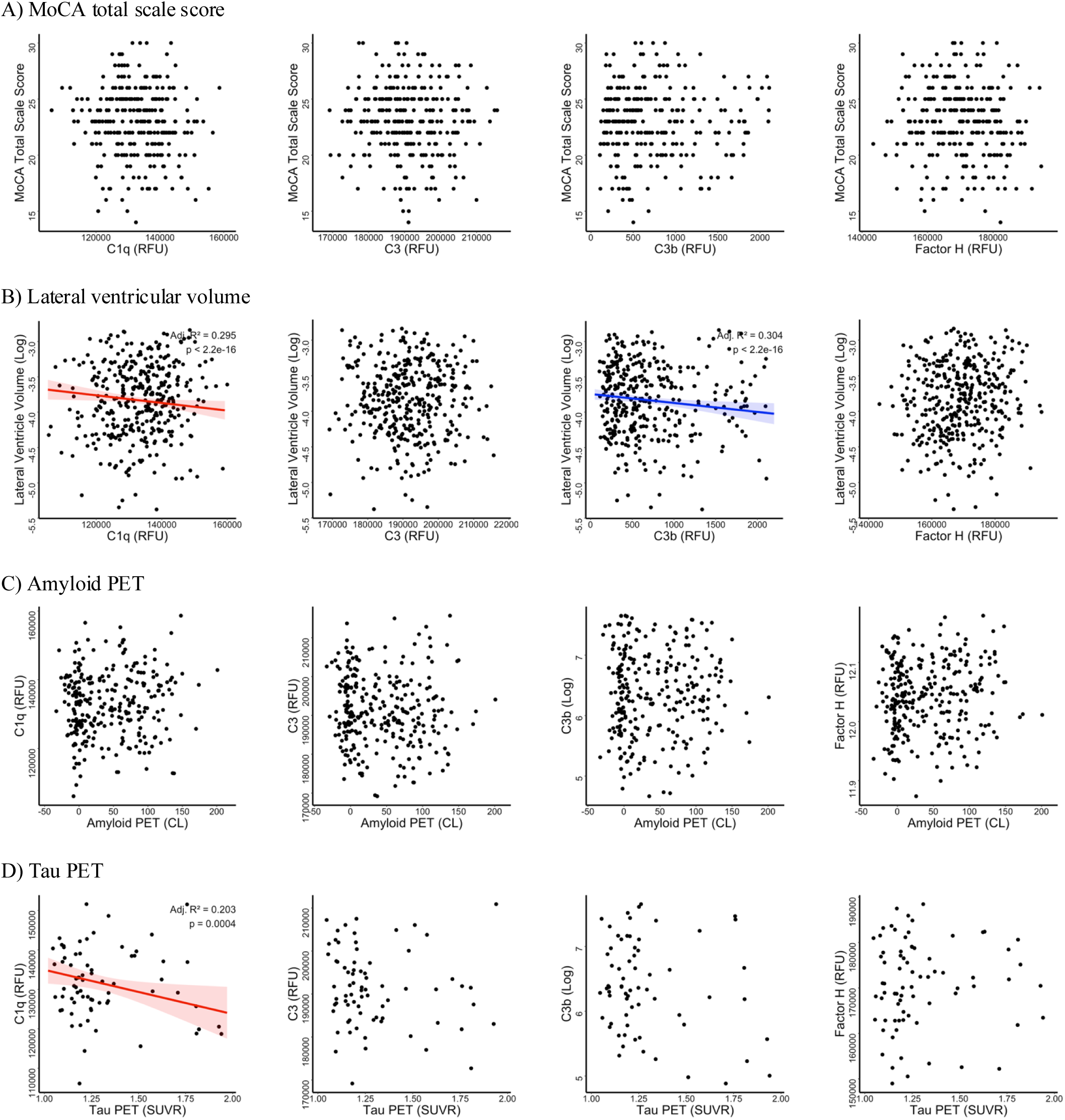
Association between baseline CSF C1q, C3, C3b, and Factor H with cognitive performance, lateral ventricular volume, and amyloid and tau PET in individuals with MCI from the ADNI cohort. CSF complement proteins C1q (n = 399), C3 (n = 397), C3b (n = 402), and Factor H (n = 396) were measured using the SomaScan 7K assay (**A**, **B**, **C**, **D**). Baseline complement protein levels were contrasted against baseline cognitive performance, assessed using the Montreal Cognitive Assessment (MoCA) (**A**, n = 291) and baseline lateral ventricle volume, measured using volumetric MRI (**B**, n = 387). Baseline amyloid PET (florbetapir; **C**, n = 287) and tau PET using data from patients’ earliest available scan (flortaucipir; **D**, n = 82) were contrasted against baseline complement protein levels. Significant linear regressions with a significant main effect of respective complement proteins or PET are indicated by a solid fitted line and shaded confidence region. Non-significant linear regression models, as well as significant models owing to main effects of covariates only (age, sex, *APOE* ε4 status, with the addition of education for MoCA score and visit year for tau PET), are depicted without fitted lines. Adjusted *R*^2^ and *P*-values are displayed in the upper right corner of each plot. *R*^2^ values are adjusted for the number of independent variables in each model, while *P*-values are corrected for multiple comparisons using the Benjamini-Hochberg (BH) method for false discovery rate (FDR) control for each dependent variable. CL = centiloids; RFU = Relative Fluorescence Units; SUVR = Standardized Uptake Value Ratio.

#### Cerebral Volumes

Left and right entorhinal cortical volumes did not show significant associations with any CSF complement protein levels in CN nor AD subjects (all *P*s *>* .05). However, left entorhinal cortical volume showed a weak negative correlation with C3 levels in MCI participants (*R*^2^ = 0.105, *P <* .0001). Regression models of left entorhinal cortical volume with C1q, C3b, and Factor H reached significance in MCI individuals only due to a negative main effect of age. Right entorhinal cortical volume also exhibited a weak negative association with C3 levels in those with MCI (*R*^2^ = 0.083, *P <* .0001). Interestingly, right entorhinal cortical volume demonstrated a weak but significant positive association with C3b levels in those with MCI (*R*^2^ = 0.100, *P <* .0001). Right entorhinal cortical volume only reached significance with C1q and Factor H in MCI subjects due to a negative main effect of age.

Regression models of left hippocampal volume against CSF complement protein levels reached significance in CN, but only owing to a negative main effect of age. In those with MCI, left hippocampal volume showed a strong and significant negative correlation with C3 levels (*R*^2^_adj_ = 0.225, *P <* .0001). Regression models of left hippocampal volume versus C1q, C3b, and Factor H levels in this group were significant only due to a negative main effect of age. Right hippocampal volume was not significantly associated with any complement protein levels in those with AD (all *P*s *>* .05), and only reached significance in those with MCI due to a negative main effect of age. Yet, right hippocampal volume showed a moderate positive correlation with C3b levels in CN subjects (*R*^2^_adj_ = 0.174, *P =* .0010). Models of right hippocampal volume versus C1q, C3, and Factor H levels in CN participants reached significance due to negative main effects of covariates. Neither left nor right hippocampal volume was significantly associated with any CSF complement protein in those with AD (all *P*s *>* .05).

Lateral ventricle volume did not show any significant correlation with any complement marker in those diagnosed with AD (all *P*s *>* .05), and only reached significance in CN subjects owing to positive main effects of covariates. Conversely, lateral ventricle volume showed strong, significant negative correlations with C1q (*R*^2^ = 0.295, *P <* .0001) and C3b (*R*^2^ = 0.304, *P <* .0001) levels in those with MCI. Associations of ventricular volume with C3 and Factor H in this group reached significance only due to negative main effects of covariates (Fig. 6B). Full cerebral volume regression statistics for CN (Table S14), MCI (Table S15), and AD (Table S16) participants are provided in the Supplementary Material.

#### Cerebral Amyloid and Tau Deposition

There was no significant relationship between amyloid PET and CSF levels of C1q, C3b, nor Factor H in CN subjects (all *P*s *>* .05). The regression model of amyloid PET versus C3 levels only reached significance due to a negative main effect of age. In those with MCI (Fig. 6C) and AD, there was no significant association between amyloid PET and C3 levels (all *P*s *>* .05), while regression models of amyloid PET versus C1q and C3b only reached significance due to main effects of covariates in both groups. Notably, amyloid PET showed a significant positive association with Factor H levels in those with AD (*R*^2^ = 0.120, *P =* .0413). This relationship was only significant in those with MCI owing to a positive main effect of male sex. Lastly, no relationship existed between tau PET and CSF complement levels in CN individuals, nor between tau PET and C3 and Factor H levels in persons with MCI (all *P*s *>* .05). However, there was a strong negative correlation between tau PET and C1q in MCI (*R*^2^ = 0.203, *p* =.0004; Fig. 6D). The association of tau PET with C3b in this group was only significant based on a main effect of male sex (Table S17).

## Discussion

Expanding on our previous work exploring novel synaptic markers as biomarkers of early tau pathology in preclinical AD,^20^ we here examined the relationship between CSF complement proteins and several domains of AD-related pathology across two complementary cohorts. In both the asymptomatic, family-history enriched PREVENT-AD cohort, and in the clinically heterogeneous ADNI cohort spanning CN, MCI, and AD individuals, the principal finding was that complement dysregulation, particularly involving C1q, is linked more consistently to tau-related pathology and synaptic injury than to cerebral amyloid burden or overt cognitive impairment. In PREVENT-AD, C1q showed robust positive associations with CSF P-tau181 and T-tau, together with several synaptic markers, while Factor H displayed similarly strong relationships with both tau and synaptic measures. In ADNI, C1q also emerged as the most consistent analyte, showing positive associations with tau biomarkers and synaptic markers across the AD spectrum. Taken together, these observations suggest that complement activation is engaged quite early in the disease process and may participate in the molecular cascade linking tau pathology to synaptic dysfunction. More importantly, these results support the view that innate immune mechanisms are not merely downstream consequences of neurodegeneration, but may be directly involved in the emergence of early synaptic vulnerability. These findings reinforce the notion that classical complement pathway activation by C1q tracks closely not just with established AD pathology,^40–43,53,54^ but with emerging tau pathology and, to a lesser degree, early amyloid processes. This pattern may also reflect compensatory upregulation of alternative complement regulators, such as Factor H, to limit excessive complement-mediated pruning and protect host cells from complement attack in the context of developing AD pathology.^55^

In contrast to the existing AD literature,^38,56,57^ C3 was not significantly related to any AD biomarkers in the asymptomatic PREVENT-AD cohort, suggesting that levels of C3 in its native inactive form may not adequately capture disease-relevant complement activity in the preclinical stage, when neuronal loss is still minimal but synaptic dysfunction already signals emerging cognitive deficits. As C3 is by and large the most abundant complement protein, it appears that complement C3 activation rather than C3 availability is more disease-relevant, as evidenced by the associations of C3b, the activated cleavage product of C3,^27^ with tau species. Interestingly, in asymptomatic PREVENT-AD subjects, CSF complement levels were not correlated with NFL, an established marker of neuronal loss. Akin to synaptic dysfunction and loss,^11,58–60^ complement activation thus seems to precede overt neurodegeneration in the pre-symptomatic phase. In further support of these findings, C3 exhibited predominantly negative associations with tau, NFL, and several synaptic markers in CN and MCI, while C3b was positively associated with NFL and multiple synaptic markers in MCI and AD in the ADNI cohort. One plausible interpretation is that total C3 and activated C3b do not index the same biological state.^61^ Total C3 may partly reflect substrate availability, turnover, or compensatory regulation, whereas C3b may better capture downstream complement activation in the context of ongoing synaptic injury.^62^ Although this study cannot resolve those mechanisms directly, the stage-dependent divergence between total C3 and C3b argues against treating all complement analytes as interchangeable indicators of inflammation.

Complement-synapse dynamics demonstrated similar patterns, wherein classical and regulatory complement proteins were widely and strongly associated with both pre-synaptic (ADAM23, GAP43, SNAP25, SYT1) and post-synaptic (ADAM22) proteins in the CSF. C1q and Factor H showed positive associations with almost all synaptic markers, with several accounting for over 30% of the variance in these complement levels. As C1q is the initiating molecule of the classical pathway, this very likely reflects its binding compromised synapses containing oligomeric tau^15,63^ to tag them for microglial elimination from both the pre- and post-synaptic side.^64^ There was again minimal association of C3 and weak association of C3b with synaptic markers, potentially signifying that C3 derivatives are prevented from over-associating with synapses to tightly control complement-driven pruning in preclinical AD. Factor H was particularly informative, but in a more complicated manner. In the pre-symptomatic PREVENT-AD cohort, Factor H showed some of the strongest associations, especially with P-tau181, T-tau, ADAM22, ADAM23, GAP43, and SYT1. By contrast, in the clinically heterogenous ADNI cohort, its associations with tau and synaptic markers in cognitively normal subjects were weaker and less consistent, while its relationship with NFL became more apparent in MCI and AD as neuronal cell loss progressively increases over time. Because Factor H is a regulatory protein of the alternative complement pathway, these findings should not be interpreted as a simple linear marker of injury. Rather, elevated Factor H in the pre-symptomatic stage may reflect an early compensatory attempt to restrain excessive complement activation in the setting of emerging tau-related synaptic stress. As disease advances, this regulatory signal may be overwhelmed and become less informative, or reflect a different balance between protection and injury.

The cognitive findings support this interpretation. In PREVENT-AD, only C1q showed a clear inverse association with global cognitive RBANS performance, whereas in ADNI, no complement protein showed an independent relationship with cognition (assessed by MoCA) after accounting for covariates. In PREVENT-AD, C3 and C3b levels correlated weakly and negatively with visuospatial ability and attention respectively, while C1q showed an inverse relationship with attention, language, and especially overall cognition measured via total RBANS score. These results could reflect the very beginning stages of cognitive decline mediated by C1q-induced synaptic alterations that precede overt clinical impairment, particularly given the high sensitivity of the RBANS relative to MoCA in detecting minute changes in cognitive ability.^50^ Rather than weakening the biological significance of the complement findings, this pattern may indicate that complement abnormalities emerge closer to molecular and synaptic dysfunction than to global clinical expression. Such a sequence would be expected in the preclinical stage of the disease, where compensatory reserve may still mask measurable cognitive consequences despite ongoing detectable pathological change. The present data therefore support the utility of complement proteins as early biological markers of disease activity, while also suggesting that their cross-sectional relationship with cognition is likely to be modest until later stages or more sensitive longitudinal analyses are considered.

In previous reports of diagnosed AD, complement levels were found to be mostly unrelated to volumes of key cerebral structures implicated in early AD, including the hippocampus and entorhinal cortex.^65,66^ While this aligns with a lack of extensive neuronal loss expected in cognitively intact individuals, C1q and Factor H levels exhibited surprising negative associations with lateral ventricular volume, such that as CSF complement levels increased, lateral ventricle measurements were smaller. The repeated inverse associations between complement markers and lateral ventricular volume in PREVENT-AD and in MCI subjects from ADNI are somewhat counterintuitive, since ventricular enlargement is usually linked to disease progression and brain atrophy. One possible interpretation is that these findings may reflect early anatomical variation such as regional glial hyperplasia typically found to be associated with glial fibrillary acidic protein (GFAP) in both CSF and plasma of MCI/AD subjects.^67^ It could also be due to selection effects within imaging subsets, or residual confounding not fully captured by the regression models. Similarly, the heterogeneous structural MRI findings for entorhinal and hippocampal volumes, particularly the discordant directions seen with C3 and C3b in ADNI, suggest that complement-related molecular changes do not map simply onto gross regional atrophy. At minimum, these results argue that CSF complement proteins may be more sensitive to active molecular and synaptic processes than to static structural outcomes. An alternative explanation could be that low-grade, complement-induced neuroinflammation may transiently alter CSF dynamics in preclinical AD stages, leading to subtle volumetric changes prior to actual atrophy. Given this unexpected directionality, these results warrant cautious interpretation and additional studies.

A notable feature of the present data is that complement proteins were related more consistently to soluble CSF AD biomarkers than to amyloid or tau PET measures, especially in PREVENT-AD. In asymptomatic, at-risk PREVENT-AD individuals, neither amyloid PET nor tau PET showed robust independent associations with complement proteins after covariate adjustment (Fig. 3C–D), despite clear relationships with CSF tau and synaptic markers. This may indicate that CSF P-tau181 indexes dynamic tau reactivity, whereas tau PET reflects years of accumulated insoluble tau burden in the brain structures. As discussed before by others,^68^ CSF and PET do not measure the same biological process, despite both indexing tau pathology. It also supports a model in which complement activity is coupled more closely to cellular stress, synaptic remodelling, and soluble tau dynamics than to plaque or tangle burden alone. The strong association between C1q and tau PET in ADNI MCI may therefore represent a later stage at which these early molecular events become detectable at the imaging level.

This study is also not without limitations. First, all analyses are cross-sectional at baseline, apart from the practical handling of ADNI tau PET timing, causality cannot be inferred. Second, the two cohorts rely on different proteomic platforms, especially Luminex/ELISA in PREVENT-AD and SomaScan in ADNI. Cross-cohort replication can thus only be interpreted at the level of direction and pattern, not absolute magnitude.

In summary, our findings support a model in which complement activation, particularly involving C1q, is engaged early in the pathobiology of AD before symptom onset, and is linked most strongly to soluble tau abnormalities and synaptic injury. It aligns complement more closely with tau biology and synaptic injury than with amyloid deposition alone, in line with emerging evidence in the field. C3b and Factor H appear to reflect related but distinct stages or regulatory components of this process, while total C3 may capture a more complex and possibly compensatory dimension of complement biology. Finally, these findings help bridge human biomarker data with the experimental literature suggesting that complement-dependent microglial pruning contributes to synapse loss. Overall, these results shift complement proteins away from being viewed as non-specific inflammatory by-products, and instead position them as stage-sensitive markers, and potentially mediators, of the tau-synapse pathway in Alzheimer’s disease. This stage-dependent framework has important implications for diagnosis and monitoring, as complement proteins may harbour potential in supplementing existing CSF biomarkers of AD. Future investigation of longitudinal complement behaviour in relation to AD progression is warranted to determine if complement levels can predict subsequent conversion to MCI and AD, rate of conversion, and severity of pathology and clinical features.

## Acknowledgements

The authors would like to thank Jennifer Tremblay-Mercier, Laurence Maligne Bruneau, Doris Dea, and Louise Théroux for their individual contributions at different stages of the project. The funding agencies played no role in the conduct of the study. JP is supported by the Fonds de recherche du Québec – Santé (FRQS) (#FRQ-356162), the Canadian Institutes of Health Research (CIHR) (PJT 153287, 178210), the Natural Sciences and Engineering Research Council of Canada (NSERC), and the J. L. Levesque Foundation. SV is supported by the FRQS (#FRQ-356162), the CIHR (PJT: 175127, 178385, 438655) and Brain Canada, whereas JCB is supported by the CIHR (PJT-175127). HZ is a Wallenberg Scholar and a Distinguished Professor at the Swedish Research Council supported by grants from the Swedish Research Council (#2023-00356, #2022-01018, #2019-02397), the European Union’s Horizon Europe research and innovation program under grant agreement No. 101053962, Swedish State Support for Clinical Research (#ALFGBG-71320), the Alzheimer Drug Discovery Foundation (ADDF), USA (#201809-2016862), the AD Strategic Fund and the Alzheimer’s Association (#ADSF-21-831376-C, #ADSF-21-831381-C, #ADSF-21-831377-C, #ADSF-24-1284328-C), the Bluefield Project, Cure Alzheimer’s Fund, the Olav Thon Foundation, the Erling-Persson Family Foundation, Familjen Rönströms Stiftelse, Stiftelsen för Gamla Tjänarinnor, Hjärnfonden, Sweden (#FO2022-0270), the European Union’s Horizon 2020 research and innovation programme under the Marie Skłodowska-Curie grant agreement No 860197 (MIRIADE), the European Union Joint Programme – Neurodegenerative Disease Research (JPND2021-00694), the National Institute for Health and Care Research University College London Hospitals Biomedical Research Centre, and the UK Dementia Research Institute at UCL (UKDRI-1003). JL is supported by the CIHR. Data used in preparation of this article were obtained from the PRe-symptomatic EValuation of Novel or Experimental Treatments for Alzheimer’s Disease (PREVENT-AD) program at the Centre for Studies on Prevention of Alzheimer’s Disease (StoP-AD), Douglas Mental Health University Institute Research Centre (http://douglas.research.mcgill.ca/stop-ad-centre). A complete listing of the PREVENT-AD Research Group can be found at: https://preventad.loris.ca/acknowledgements/acknowledgements.php?date=2026-05-15.

## Data Availability

The multimodal data, including subject characteristics, CSF biomarker levels, tau and amyloid PET SUVR, *APOE* genotypes, and RBANS scores, have been summarized in the PREVENT-AD Data Release 7.0, available from the corresponding authors upon reasonable request.

## Author Contributions

JL, JP, CP, SV, and JB conceptualized the research. JL, CP, MS, and HZ performed CSF biomarkers measurements, data quality control, data compilation whereas SD performed lumbar punctures in the PREVENT-AD participants. JL, JP, CP, MS, SV, and JB contributed to data analysis. JL, JP, CP, MS, and SV developed the algorithms for data analysis. JL, JP, JB and SV wrote the original manuscript draft. All authors reviewed, edited, and approved the final manuscript.

