## Supplementary Material for "CSF complement proteins are associated with early tau pathology and synaptic damage in an asymptomatic population at risk of Alzheimer’s disease"

***Table S1***. Demographics for cognitively normal (CN) individuals in the ADNI cohort.

| n = 167 | Sex | | Sig. | *APOE* ε4 Status | | Sig. |
| --- | --- | --- | --- | --- | --- | --- |
|  | **Male  (n = 89)** | **Female  (n = 78)** |  | **ε4– (n = 126)** | **ε4+ (n = 41)** |  |
| Age (years) | 75.19 ± 6.22 | 73.54 ± 5.50 |  | 74.50 ± 5.96 | 74.15 ± 5.94 |  |
| CSF Aβ42 (RFU) | 1042.84 ± 377.45 | 1048.02 ± 361.10 |  | 1130.32 ± 357.36 | 848.92 ± 319.35 | *** |
| CSF P-tau181 (RFU) | 20.91 ± 7.28 | 20.81 ± 7.32 |  | 20.03 ± 6.96 | 23.61 ± 7.72 | * |
| CSF T-tau (RFU) | 229.63 ± 75.08 | 228.73 ± 75.93 |  | 223.56 ± 75.42 | 248.14 ± 72.45 |  |
| CSF NFL (RFU) | 987.73 ± 222.03 | 852.06 ± 204.39 | *** | 938.43 ± 225.12 | 879.41 ± 216.45 |  |
| CSF C1q (RFU) | 135068.95 ± 9331.71 | 132734.19 ± 8460.80 |  | 134180.04 ± 8783.58 | 133414.24 ± 9720.57 |  |
| CSF C3 (RFU) | 192379.65 ± 8637.28 | 195024.27 ± 8390.16 |  | 193196.30 ± 8524.68 | 194856.6 ± 8814.13 |  |
| CSF C3b (RFU) | 604.72 ± 301.68 | 527.98 ± 303.32 |  | 564.62 ± 295.21 | 583.70 ± 334.52 |  |
| CSF Factor H (RFU) | 168434.24 ± 9091.10 | 164817.40 ± 9945.41 | * | 167041.31 ± 9549.85 | 165915.14 ± 9974.87 |  |
| CSF ADAM22 (RFU) | 43146.27 ± 9359.20 | 41592.33 ± 11263.04 |  | 42259.09 ± 10263.94 | 42911.06 ± 10476.15 |  |
| CSF ADAM23 (RFU) | 4107.91 ± 1079.22 | 4052.96 ± 1041.34 |  | 4074.13 ± 1056.38 | 4110.40 ± 1081.44 |  |
| CSF GAP43 (RFU) | 1845.17 ± 539.84 | 1839.62 ± 561.68 |  | 1817.39 ± 554.40 | 1924.26 ± 526.74 |  |
| CSF SNAP25 (RFU) | 110.91 ± 5.81 | 111.10 ± 6.11 |  | 111.19 ± 5.87 | 110.44 ± 6.16 |  |
| CSF SYT1 (RFU) | 4417.61 ± 1096.51 | 4497.21 ± 1225.99 |  | 4393.74 ± 1128.69 | 4639.29 ± 1230.27 |  |
| MRI volume – entorhinal cortex (mm^3^) | 3998.94 ± 517.60 | 3722.39 ± 566.75 |  | 3887.38 ± 580.05 | 3803.62 ± 473.87 |  |
| MRI volume – hippocampus (mm^3^) | 7580.46 ± 759.74 | 7164.84 ± 766.12 | ** | 7367.99 ± 787.68 | 7442.06 ± 798.95 |  |
| MRI volume – lateral ventricles (mm^3^) | 34655.61 ± 12339.05 | 27205.01 ± 13831.47 | ** | 30793.47 ± 14004.06 | 31769.94 ± 12258.76 |  |
| PET amyloid (CL) | 6.26 ± 22.63 | 9.13 ± 21.84 |  | 7.24 ± 22.05 | 9.50 ± 23.49 |  |
| PET tau (SUVR) | 1.20 ± 0.09 | 1.20 ± 0.08 |  | 1.19 ± 0.08 | 1.24 ± 0.09 | * |
| MoCA – total score | 25.55 ± 2.15 | 26.12 ± 2.54 |  | 25.99 ± 2.36 | 25.26 ± 2.32 |  |

*APOE* apolipoprotein E, *CSF* cerebrospinal fluid, *Aβ42* amyloid-beta 42, *P-tau181* phosphorylated tau 181, *T-tau* total tau, *NFL* neurofilament light, *RFU* relative fluorescence units, *ADAM* a disintegrin and metalloproteinase domain-containing protein, *GAP43* growth-associated protein 43, *SNAP25* synaptosomal-associated protein of 25 kDa, *SYT1* synatotagmin-1, *MRI* magnetic resonance imaging, *PET* positron emission tomography, *CL* centiloids, *MoCA* Montreal Cognitive Assessment.

* = *P* < 0.05; ** = *P* < 0.01; *** = *P* < 0.001; **** = *P* < .0001

***Table S2***. Demographics for individuals with mild cognitive impairment (MCI) in the ADNI cohort.

| n = 403 | Sex | | Sig. | *APOE* ε4 Status | | Sig. |
| --- | --- | --- | --- | --- | --- | --- |
|  | **Male  (n = 232)** | **Female  (n = 171)** |  | **ε4– (n = 193)** | **ε4+  (n = 210)** |  |
| Age (years) | 73.41 ± 7.33 | 70.76 ± 7.59 |  | 73.02 ± 7.96 | 71.60 ± 7.10 |  |
| CSF Aβ42 (RFU) | 825.28 ± 344.68 | 881.52 ± 339.75 |  | 996.83 ± 346.05 | 735.96 ± 295.72 |  |
| CSF P-tau181 (RFU) | 25.63 ± 11.82 | 27.18 ± 12.58 |  | 21.50 ± 10.10 | 30.75 ± 12.22 | **** |
| CSF T-tau (RFU) | 268.75 ± 111.05 | 283.71 ± 112.61 |  | 232.27 ± 93.29 | 314.45 ± 113.21 | **** |
| CSF NFL (RFU) | 1060.71 ± 294.55 | 908.51 ± 286.03 | **** | 1000.57 ± 299.01 | 990.62 ± 301.88 | **** |
| CSF C1q (RFU) | 134726.44 ± 9554.04 | 131573.29 ± 10015.63 | ** | 134198.82 ± 9514.80 | 132716.83 ± 10130.58 |  |
| CSF C3 (RFU) | 190553.98 ± 9030.89 | 192802.99 ± 9302.10 | * | 191442.11 ± 8827.92 | 191558.07 ± 9555.18 |  |
| CSF C3b (RFU) | 782.24 ± 510.93 | 578.53 ± 409.48 | **** | 777.36 ± 529.02 | 613.35 ± 412.16 |  |
| CSF Factor H (RFU) | 171022.68 ± 9544.69 | 166920.48 ± 9219.78 | **** | 169143.16 ± 9414.66 | 169409.44 ± 9812.04 | *** |
| CSF ADAM22 (RFU) | 40451.38 ± 9020.83 | 40623.26 ± 9219.46 |  | 39835.98 ± 9117.83 | 41148.00 ± 9053.36 |  |
| CSF ADAM23 (RFU) | 3930.46 ± 1113.54 | 4248.66 ± 1181.69 | * | 3921.83 ± 1129.74 | 4195.78 ± 1159.46 |  |
| CSF GAP43 (RFU) | 1972.56 ± 688.73 | 2026.13 ± 608.36 |  | 1838.93 ± 616.43 | 2143.70 ± 662.29 | * |
| CSF SNAP25 (RFU) | 110.88 ± 5.54 | 109.63 ± 5.96 |  | 110.69 ± 5.62 | 110.03 ± 5.86 | **** |
| CSF SYT1 (RFU) | 4401.33 ± 1272.44 | 4678.94 ± 1296.09 | * | 4249.51 ± 1231.11 | 4765.47 ± 1292.83 |  |
| MRI volume – entorhinal cortex (mm^3^) | 3681.69 ± 754.75 | 3331.54 ± 621.66 |  | 3580.84 ± 689.46 | 3482.06 ± 745.92 | *** |
| MRI volume – hippocampus (mm^3^) | 6967.39 ± 1205.98 | 6609.07 ± 1086.34 | *** | 6915.53 ± 1184.95 | 6709.22 ± 1143.91 |  |
| MRI volume – lateral ventricles (mm^3^) | 42118.69 ± 19180.18 | 29592.41 ± 15225.88 | **** | 38704.42 ± 19034.75 | 34788.69 ± 18067.35 |  |
| PET amyloid (CL) | 44.78 ± 48.32 | 41.13 ± 48.61 |  | 20.46 ± 39.37 | 64.84 ± 46.32 | * |
| PET tau (SUVR) | 1.26 ± 0.19 | 1.28 ± 0.20 |  | 1.20 ± 0.13 | 1.36 ± 0.23 | **** |
| MoCA – total score | 23.04 ± 2.98 | 23.18 ± 3.22 |  | 23.50 ± 3.07 | 22.70 ± 3.04 | **** |

*APOE* apolipoprotein E, *CSF* cerebrospinal fluid, *Aβ42* amyloid-beta 42, *P-tau181* phosphorylated tau 181, *T-tau* total tau, *NFL* neurofilament light, *RFU* relative fluorescence units, *ADAM* a disintegrin and metalloproteinase domain-containing protein, *GAP43* growth-associated protein 43, *SNAP25* synaptosomal-associated protein of 25 kDa, *SYT1* synatotagmin-1, *MRI* magnetic resonance imaging, *PET* positron emission tomography, *CL* centiloids, *MoCA* Montreal Cognitive Assessment.

* = *P* < 0.05; ** = *P* < 0.01; *** = *P* < 0.001; **** = *P* < .0001

***Table S3***. Demographics for individuals with Alzheimer’s disease (AD) in the ADNI cohort.

| n = 138 | Sex | | Sig. | *APOE* ε4 Status | | Sig. |
| --- | --- | --- | --- | --- | --- | --- |
|  | **Male  (n = 84)** | **Female  (n = 54)** |  | **ε4– (n = 43)** | **ε4+  (n = 95)** |  |
| Age (years) | 76.04 ± 8.07 | 74.28 ± 9.17 |  | 78.05 ± 9.09 | 74.13 ± 8.02 |  |
| CSF Aβ42 (RFU) | 581.71 ± 198.97 | 600.61 ± 170.71 |  | 648.76 ± 232.92 | 567.00 ± 164.00 |  |
| CSF P-tau181 (RFU) | 35.67 ± 14.58 | 36.16 ± 13.44 |  | 35.39 ± 18.08 | 36.09 ± 11.88 |  |
| CSF T-tau (RFU) | 353.13 ± 128.83 | 372.84 ± 136.91 |  | 359.69 ± 161.15 | 361.41 ± 116.77 |  |
| CSF NFL (RFU) | 1192.33 ± 284.32 | 1054.98 ± 292.17 | * | 1242.18 ± 331.71 | 1098.17 ± 269.19 | * |
| CSF C1q (RFU) | 133360.36 ± 8903.11 | 131501.19 ± 9882.29 |  | 135266.22 ± 10140.62 | 131419.63 ± 8682.71 |  |
| CSF C3 (RFU) | 189043.77 ± 6159.91 | 193829.64 ± 10154.67 | ** | 191258.53 ± 8338.30 | 190747.18 ± 8266.34 |  |
| CSF C3b (RFU) | 695.44 ± 416.20 | 493.98 ± 362.35 | ** | 633.79 ± 388.77 | 602.49 ± 414.73 |  |
| CSF Factor H (RFU) | 171677.58 ± 9058.68 | 167339.10 ± 10991.09 | * | 173337.48 ± 10004.91 | 168489.75 ± 9745.85 | * |
| CSF ADAM22 (RFU) | 40818.93 ± 8860.51 | 37816.28 ± 8026.35 |  | 40255.65 ± 10335.36 | 39385.05 ± 7776.31 |  |
| CSF ADAM23 (RFU) | 3804.02 ± 1044.77 | 3683.04 ± 1018.18 |  | 3903.54 ± 1108.52 | 3691.34 ± 995.68 |  |
| CSF GAP43 (RFU) | 2187.57 ± 693.03 | 2151.73 ± 688.80 |  | 2191.41 ± 787.83 | 2165.56 ± 643.43 |  |
| CSF SNAP25 (RFU) | 112.77 ± 6.52 | 111.12 ± 6.33 |  | 112.24 ± 6.42 | 112.07 ± 6.53 |  |
| CSF SYT1 (RFU) | 4549.54 ± 1152.47 | 4410.29 ± 1244.03 |  | 4482.60 ± 1347.80 | 4501.19 ± 1115.38 |  |
| MRI volume – entorhinal cortex (mm^3^) | 3010.14 ± 682.39 | 2686.53 ± 579.14 |  | 3063.86 ± 735.12 | 2795.20 ± 612.91 | * |
| MRI volume – hippocampus (mm^3^) | 6057.50 ± 920.53 | 5662.71 ± 1005.32 |  | 6060.00 ± 1155.72 | 5831.54 ± 878.12 |  |
| MRI volume – lateral ventricles (mm^3^) | 48946.17 ± 19302.96 | 38206.40 ± 12728.11 |  | 43487.36 ± 16123.34 | 44940.62 ± 18339.55 |  |
| PET amyloid (CL) | 92.10 ± 38.09 | 102.69 ± 23.69 |  | 93.00 ± 38.84 | 97.41 ± 31.62 |  |
| MoCA – total score | 17.65 ± 4.02 | 16.24 ± 4.78 |  | 16.00 ± 4.14 | 17.57 ± 4.37 |  |

*APOE* apolipoprotein E, *CSF* cerebrospinal fluid, *Aβ42* amyloid-beta 42, *P-tau181* phosphorylated tau 181, *T-tau* total tau, *NFL* neurofilament light, *RFU* relative fluorescence units, *ADAM* a disintegrin and metalloproteinase domain-containing protein, *GAP43* growth-associated protein 43, *SNAP25* synaptosomal-associated protein of 25 kDa, *SYT1* synatotagmin-1, *MRI* magnetic resonance imaging, *PET* positron emission tomography, *CL* centiloids, *MoCA* Montreal Cognitive Assessment.

* = *P* < 0.05; ** = *P* < 0.01; *** = *P* < 0.001; **** = *P* < .0001

***Table S4*.** Summary of RBANS index score regression analyses in the PREVENT-AD cohort. All analyses were adjusted for participant age, sex, *APOE* ε4 genotype, years of education, and test version. Significant models with a significant main effect of complement protein are highlighted in yellow.

| **RBANS score** | **Complement protein** | **Adjusted R^2^** | **Main effect of complement protein** | **Significance level (FDR)** |
| --- | --- | --- | --- | --- |
| **Attention** | C1q | 0.088 | Negative | .0034 |
|  | C3 | 0.043 | None | .0497 |
|  | C3b | 0.080 | Negative | .0082 |
|  | Factor H | 0.049 | None | .0351 |
| **Delayed memory** | C1q | 0 | – | n.s. |
|  | C3 | 0.001 | – | n.s. |
|  | C3b | 0.002 | – | n.s. |
|  | Factor H | 0.010 | – | n.s. |
| **Immediate memory** | C1q | 0.106 | None | .0016 |
|  | C3 | 0.084 | None | .0056 |
|  | C3b | 0.104 | None | .0029 |
|  | Factor H | 0.091 | None | .0033 |
| **Language** | C1q | 0.128 | Negative | .0002 |
|  | C3 | 0.093 | None | .0015 |
|  | C3b | 0.097 | None | .0006 |
|  | Factor H | 0.096 | None | .0016 |
| **Visuospatial constructional** | C1q | 0.073 | None | .0077 |
|  | C3 | 0.105 | Positive | .0015 |
|  | C3b | 0.049 | None | .0412 |
|  | Factor H | 0.094 | None | .0029 |
| **Total scale** | C1q | 0.183 | Negative | < .0001 |
|  | C3 | 0.137 | None | .0004 |
|  | C3b | 0.142 | None | .0005 |
|  | Factor H | 0.151 | None | .0001 |

***Table S5*.** Summary of ICV-corrected brain volume regression analyses in the PREVENT-AD cohort. All analyses were adjusted for participant age, sex, and *APOE* ε4 genotype. Significant models with a significant main effect of complement protein are highlighted in yellow.

| **Brain region** | **Complement protein** | **Adjusted R^2^** | **Main effect of complement protein** | **Significance level (FDR)** |
| --- | --- | --- | --- | --- |
| **Left entorhinal cortex** | C1q | 0.004 | – | n.s. |
|  | C3 | 0.018 | – | n.s. |
|  | C3b | 0.020 | – | n.s. |
|  | Factor H | 0.015 | – | n.s. |
| **Right entorhinal cortex** | C1q | 0 | – | n.s. |
|  | C3 | 0 | – | n.s. |
|  | C3b | 0.002 | – | n.s. |
|  | Factor H | 0 | – | n.s. |
| **Left hippocampus** | C1q | 0.127 | None | .0007 |
|  | C3 | 0.112 | None | .0018 |
|  | C3b | 0.112 | None | .0019 |
|  | Factor H | 0.133 | None | .0004 |
| **Right hippocampus** | C1q | 0.131 | None | .0007 |
|  | C3 | 0.127 | None | .0011 |
|  | C3b | 0.125 | None | .0013 |
|  | Factor H | 0.134 | None | .0004 |
| **Lateral ventricles** | C1q | 0.256 | Negative | < .0001 |
|  | C3 | 0.239 | None | < .0001 |
|  | C3b | 0.231 | None | < .0001 |
|  | Factor H | 0.281 | Negative | < .0001 |

***Table S6*.** Summary of amyloid and tau PET regression analyses in the PREVENT-AD cohort. All analyses were adjusted for participant age, sex, and *APOE* ε4 genotype. Significant models with a significant main effect of complement protein are highlighted in yellow.

| **Brain region** | **Complement protein** | **Adjusted R^2^** | **Main effect of complement protein** | **Significance level (FDR)** |
| --- | --- | --- | --- | --- |
| **Amyloid – total index** | C1q | 0.149 | None | .0045 |
|  | C3 | 0 | – | n.s. |
|  | C3b | 0 | – | n.s. |
|  | Factor H | 0.148 | None | .0045 |
| **Tau – entorhinal cortex** | C1q | 0.099 | None | .0231 |
|  | C3 | 0 | – | n.s. |
|  | C3b | 0 | – | n.s. |
|  | Factor H | 0.166 | None | .0019 |
| **Tau – fusiform gyrus** | C1q | 0.103 | None | .0182 |
|  | C3 | 0 | – | n.s. |
|  | C3b | 0.011 | – | n.s. |
|  | Factor H | 0.165 | None | .0017 |
| **Tau – lingual gyrus** | C1q | 0.107 | None | .0153 |
|  | C3 | 0 | – | n.s. |
|  | C3b | 0.030 | – | n.s. |
|  | Factor H | 0.177 | None | .0009 |
| **Tau – metaROI** | C1q | 0.105 | None | .0165 |
|  | C3 | 0 | – | n.s. |
|  | C3b | 0.016 | – | n.s. |
|  | Factor H | 0.168 | None | .0015 |

***Table S7*.** Summary of AD biomarker regression analyses in cognitively normal (CN) individuals in the ADNI cohort. All analyses were adjusted for participant age, sex, and *APOE* ε4 genotype. Significant models with a significant main effect of AD biomarker are highlighted in yellow.

| **Complement protein** | **AD biomarker** | **Adjusted R^2^** | **Main effect of biomarker** | **Significance level (FDR)** |
| --- | --- | --- | --- | --- |
| **C1q** | Aβ42 | 0.051 | – | n.s. |
|  | P*tau*-181 | 0.089 | Positive | .0026 |
|  | T*tau* | 0.100 | Positive | .0011 |
|  | NFL | 0.106 | Positive | .0012 |
| **C3** | Aβ42 | 0.114 | Negative | .0054 |
|  | P*tau*-181 | 0.087 | Negative | .0026 |
|  | T*tau* | 0.098 | Negative | .0011 |
|  | NFL | 0.072 | Negative | .0043 |
| **C3b** | Aβ42 | 0 | – | n.s. |
|  | P*tau*-181 | 0.027 | – | n.s. |
|  | T*tau* | 0.034 | – | n.s. |
|  | NFL | 0.099 | Positive | .0042 |
| **Factor H** | Aβ42 | 0.028 | – | n.s. |
|  | P*tau*-181 | 0.033 | – | n.s. |
|  | T*tau* | 0.034 | – | n.s. |
|  | NFL | 0.063 | Positive | .0066 |

***Table S8*.** Summary of AD biomarker regression analyses in individuals with mild cognitive impairment (MCI) in the ADNI cohort. All analyses were adjusted for participant age, sex, and *APOE* ε4 genotype. Significant models with a significant main effect of AD biomarker are highlighted in yellow.

| **Complement protein** | **AD biomarker** | **Adjusted R^2^** | **Main effect of biomarker** | **Significance level (FDR)** |
| --- | --- | --- | --- | --- |
| **C1q** | Aβ42 | 0.098 | None | < .0001 |
|  | P*tau*-181 | 0.133 | Positive | < .0001 |
|  | T*tau* | 0.137 | Positive | < .0001 |
|  | NFL | 0.125 | Positive | < .0001 |
| **C3** | Aβ42 | 0.031 | Negative | .0056 |
|  | P*tau*-181 | 0.041 | Negative | .0006 |
|  | T*tau* | 0.051 | Negative | < .0001 |
|  | NFL | 0.043 | Negative | .0004 |
| **C3b** | Aβ42 | 0.060 | None | .0002 |
|  | P*tau*-181 | 0.087 | Positive | < .0001 |
|  | T*tau* | 0.096 | Positive | < .0001 |
|  | NFL | 0.119 | Positive | < .0001 |
| **Factor H** | Aβ42 | 0.041 | None | .0014 |
|  | P*tau*-181 | 0.065 | None | < .0001 |
|  | T*tau* | 0.066 | None | < .0001 |
|  | NFL | 0.057 | Positive | < .0001 |

***Table S9*.** Summary of AD biomarker regression analyses in individuals with Alzheimer’s disease (AD) in the ADNI cohort. All analyses were adjusted for participant age, sex, and *APOE* ε4 genotype. Significant models with a significant main effect of AD biomarker are highlighted in yellow.

| **Complement protein** | **AD biomarker** | **Adjusted R^2^** | **Main effect of biomarker** | **Significance level (FDR)** |
| --- | --- | --- | --- | --- |
| **C1q** | Aβ42 | 0.149 | Positive | .0003 |
|  | P*tau*-181 | 0.157 | Positive | .0001 |
|  | T*tau* | 0.165 | Positive | < .0001 |
|  | NFL | 0.164 | Positive | < .0001 |
| **C3** | Aβ42 | 0.101 | None | .0014 |
|  | P*tau*-181 | 0.096 | None | .0013 |
|  | T*tau* | 0.098 | None | .0012 |
|  | NFL | 0.133 | None | .0003 |
| **C3b** | Aβ42 | 0.062 | None | .0222 |
|  | P*tau*-181 | 0.103 | None | .0019 |
|  | T*tau* | 0.106 | None | .0016 |
|  | NFL | 0.285 | Positive | < .0001 |
| **Factor H** | Aβ42 | 0.085 | None | .0064 |
|  | P*tau*-181 | 0.108 | None | .0013 |
|  | T*tau* | 0.111 | None | .0012 |
|  | NFL | 0.133 | Positive | .0003 |

***Table S10*.** Summary of synaptic marker regression analyses in cognitively normal (CN) individuals in the ADNI cohort. All analyses were adjusted for participant age, sex, and *APOE* ε4 genotype. Significant models with a significant main effect of synaptic marker are highlighted in yellow.

| **Complement protein** | **Synaptic marker** | **Adjusted R^2^** | **Main effect of synaptic marker** | **Significance level (FDR)** |
| --- | --- | --- | --- | --- |
| **C1q** | ADAM22 | 0.151 | Positive | < .0001 |
|  | ADAM23 | 0.122 | Positive | .0005 |
|  | GAP43 | 0.110 | Positive | .0023 |
|  | SNAP25 | 0.028 | – | n.s. |
|  | SYT1 | 0.116 | Positive | .0009 |
| **C3** | ADAM22 | 0.096 | Negative | .0010 |
|  | ADAM23 | 0.082 | Negative | .0036 |
|  | GAP43 | 0.087 | Negative | .0023 |
|  | SNAP25 | 0.040 | – | n.s. |
|  | SYT1 | 0.096 | Negative | .0009 |
| **C3b** | ADAM22 | 0.025 | – | n.s. |
|  | ADAM23 | 0.037 | – | n.s. |
|  | GAP43 | 0.029 | – | n.s. |
|  | SNAP25 | 0.001 | – | n.s. |
|  | SYT1 | 0.026 | – | n.s. |
| **Factor H** | ADAM22 | 0.019 | – | n.s. |
|  | ADAM23 | 0.036 | – | n.s. |
|  | GAP43 | 0.030 | – | n.s. |
|  | SNAP25 | 0.011 | – | n.s. |
|  | SYT1 | 0.026 | – | n.s. |

***Table S11*.** Summary of synaptic marker regression analyses in individuals with mild cognitive impairment (MCI) in the ADNI cohort. All analyses were adjusted for participant age, sex, and *APOE* ε4 genotype. Significant models with a significant main effect of synaptic marker are highlighted in yellow.

| **Complement protein** | **Synaptic marker** | **Adjusted R^2^** | **Main effect of synaptic marker** | **Significance level (FDR)** |
| --- | --- | --- | --- | --- |
| **C1q** | ADAM22 | 0.139 | Positive | < .0001 |
|  | ADAM23 | 0.149 | Positive | < .0001 |
|  | GAP43 | 0.147 | Positive | < .0001 |
|  | SNAP25 | 0.086 | None | < .0001 |
|  | SYT1 | 0.157 | Positive | < .0001 |
| **C3** | ADAM22 | 0.072 | Negative | < .0001 |
|  | ADAM23 | 0.080 | Negative | < .0001 |
|  | GAP43 | 0.053 | Negative | < .0001 |
|  | SNAP25 | 0.007 | – | n.s. |
|  | SYT1 | 0.071 | Negative | < .0001 |
| **C3b** | ADAM22 | 0.096 | Positive | < .0001 |
|  | ADAM23 | 0.126 | Positive | < .0001 |
|  | GAP43 | 0.093 | Positive | < .0001 |
|  | SNAP25 | 0.057 | None | .0001 |
|  | SYT1 | 0.094 | Positive | < .0001 |
| **Factor H** | ADAM22 | 0.058 | Positive | < .0001 |
|  | ADAM23 | 0.075 | Positive | < .0001 |
|  | GAP43 | 0.069 | None | < .0001 |
|  | SNAP25 | 0.047 | None | .0003 |
|  | SYT1 | 0.052 | None | < .0001 |

***Table S12*.** Summary of synaptic marker regression analyses in individuals with Alzheimer’s disease (AD) in the ADNI cohort. All analyses were adjusted for participant age, sex, and *APOE* ε4 genotype. Significant models with a significant main effect of synaptic marker are highlighted in yellow.

| **Complement protein** | **Synaptic marker** | **Adjusted R^2^** | **Main effect of synaptic marker** | **Significance level (FDR)** |
| --- | --- | --- | --- | --- |
| **C1q** | ADAM22 | 0.177 | Positive | < .0001 |
|  | ADAM23 | 0.196 | Positive | < .0001 |
|  | GAP43 | 0.201 | Positive | < .0001 |
|  | SNAP25 | 0.124 | None | .0005 |
|  | SYT1 | 0.143 | Positive | .0004 |
| **C3** | ADAM22 | 0.105 | Negative | .0011 |
|  | ADAM23 | 0.101 | None | .0019 |
|  | GAP43 | 0.097 | None | .0019 |
|  | SNAP25 | 0.097 | Negative | .0006 |
|  | SYT1 | 0.110 | None | .0012 |
| **C3b** | ADAM22 | 0.132 | Positive | .0012 |
|  | ADAM23 | 0.106 | None | .0019 |
|  | GAP43 | 0.114 | None | .0019 |
|  | SNAP25 | 0.169 | Positive | .0001 |
|  | SYT1 | 0.113 | Positive | .0029 |
| **Factor H** | ADAM22 | 0.100 | None | .0015 |
|  | ADAM23 | 0.096 | None | .0019 |
|  | GAP43 | 0.101 | None | .0019 |
|  | SNAP25 | 0.135 | None | .0003 |
|  | SYT1 | 0.097 | None | .0026 |

***Table S13*.** Summary of MoCA total score regression analyses in the ADNI cohort. All analyses were adjusted for participant age, sex, *APOE* ε4 genotype, and years of education. Significant models with a significant main effect of complement protein are highlighted in yellow.

| **Diagnosis** | **Complement protein** | **Adjusted R^2^** | **Main effect of complement protein** | **Significance level (FDR)** |
| --- | --- | --- | --- | --- |
| **Cognitively normal (CN)** | C1q | 0.091 | None | .0205 |
|  | C3 | 0.117 | None | .0165 |
|  | C3b | 0.072 | None | .0389 |
|  | Factor H | 0.090 | None | .0205 |
| **Mild cognitive impairment (MCI)** | C1q | 0.092 | None | < .0001 |
|  | C3 | 0.102 | None | < .0001 |
|  | C3b | 0.125 | None | < .0001 |
|  | Factor H | 0.106 | None | < .0001 |
| **Alzheimer’s disease (AD)** | C1q | 0.134 | None | .0013 |
|  | C3 | 0.116 | None | .0092 |
|  | C3b | 0.130 | None | .0092 |
|  | Factor H | 0.116 | None | .0092 |

***Table S14*.** Summary of ICV-corrected brain volume regression analyses in cognitively normal (CN) individuals in the ADNI cohort. All analyses were adjusted for participant age, sex, and *APOE* ε4 genotype. Significant models with a significant main effect of complement protein are highlighted in yellow.

| **Brain region** | **Complement protein** | **Adjusted R^2^** | **Main effect of complement protein** | **Significance level (FDR)** |
| --- | --- | --- | --- | --- |
| **Left entorhinal cortex** | C1q | 0 | – | n.s. |
|  | C3 | .006 | – | n.s. |
|  | C3b | 0 | – | n.s. |
|  | Factor H | 0.011 | – | n.s. |
| **Right entorhinal cortex** | C1q | 0 | – | n.s. |
|  | C3 | 0 | – | n.s. |
|  | C3b | 0 | – | n.s. |
|  | Factor H | 0 | – | n.s. |
| **Left hippocampus** | C1q | 0.088 | None | .0241 |
|  | C3 | 0.094 | None | .0179 |
|  | C3b | 0.083 | None | .0372 |
|  | Factor H | 0.120 | None | .0055 |
| **Right hippocampus** | C1q | 0.173 | None | .0006 |
|  | C3 | 0.179 | None | .0004 |
|  | C3b | 0.174 | Positive | .0010 |
|  | Factor H | 0.187 | None | .0003 |
| **Lateral ventricles** | C1q | 0.317 | None | < .0001 |
|  | C3 | 0.317 | None | < .0001 |
|  | C3b | 0.277 | None | < .0001 |
|  | Factor H | 0.322 | None | < .0001 |

***Table S15*.** Summary of ICV-corrected brain volume regression analyses in individuals with mild cognitive impairment (MCI) in the ADNI cohort. All analyses were adjusted for participant age, sex, and *APOE* ε4 genotype. Significant models with a significant main effect of complement protein are highlighted in yellow.

| **Brain region** | **Complement protein** | **Adjusted R^2^** | **Main effect of complement protein** | **Significance level (FDR)** |
| --- | --- | --- | --- | --- |
| **Left entorhinal cortex** | C1q | 0.077 | None | < .0001 |
|  | C3 | 0.105 | Negative | < .0001 |
|  | C3b | 0.087 | None | < .0001 |
|  | Factor H | 0.086 | None | < .0001 |
| **Right entorhinal cortex** | C1q | 0.070 | None | .0001 |
|  | C3 | 0.083 | Negative | < .0001 |
|  | C3b | 0.100 | Positive | < .0001 |
|  | Factor H | 0.077 | None | < .0001 |
| **Left hippocampus** | C1q | 0.209 | None | < .0001 |
|  | C3 | 0.225 | Negative | < .0001 |
|  | C3b | 0.227 | None | < .0001 |
|  | Factor H | 0.213 | None | < .0001 |
| **Right hippocampus** | C1q | 0.206 | None | < .0001 |
|  | C3 | 0.205 | None | < .0001 |
|  | C3b | 0.217 | None | < .0001 |
|  | Factor H | 0.205 | None | < .0001 |
| **Lateral ventricles** | C1q | 0.295 | Negative | < .0001 |
|  | C3 | 0.293 | None | < .0001 |
|  | C3b | 0.304 | Negative | < .0001 |
|  | Factor H | 0.291 | None | < .0001 |

***Table S16*.** Summary of ICV-corrected brain volume regression analyses in individuals with Alzheimer’s disease (AD) in the ADNI cohort. All analyses were adjusted for participant age, sex, and *APOE* ε4 genotype. Significant models with a significant main effect of AD biomarker are highlighted in yellow.

| **Brain region** | **Complement protein** | **Adjusted R^2^** | **Main effect of complement protein** | **Significance level (FDR)** |
| --- | --- | --- | --- | --- |
| **Left entorhinal cortex** | C1q | 0.055 | – | n.s. |
|  | C3 | 0.057 | – | n.s. |
|  | C3b | 0 | – | n.s. |
|  | Factor H | 0.058 | – | n.s. |
| **Right entorhinal cortex** | C1q | 0 | – | n.s. |
|  | C3 | 0 | – | n.s. |
|  | C3b | 0 | – | n.s. |
|  | Factor H | 0.009 | – | n.s. |
| **Left hippocampus** | C1q | 0.133 | – | n.s. |
|  | C3 | 0.094 | – | n.s. |
|  | C3b | 0.104 | – | n.s. |
|  | Factor H | 0.103 | – | n.s. |
| **Right hippocampus** | C1q | 0.077 | – | n.s. |
|  | C3 | 0.063 | – | n.s. |
|  | C3b | 0.073 | – | n.s. |
|  | Factor H | 0.061 | – | n.s. |
| **Lateral ventricles** | C1q | 0 | – | n.s. |
|  | C3 | 0 | – | n.s. |
|  | C3b | 0.010 | – | n.s. |
|  | Factor H | 0.011 | – | n.s. |

***Table S17*.** Summary of PET regression analyses in the ADNI cohort. All analyses were adjusted for participant age, sex, and *APOE* ε4 genotype, as well as visit year for tau PET analyses. Significant models with a significant main effect of complement protein are highlighted in yellow.

| **Diagnosis** | **AD PET biomarker** | **Complement protein** | **Adjusted R^2^** | **Main effect of complement protein** | **Significance level (FDR)** |
| --- | --- | --- | --- | --- | --- |
| **Cognitively normal (CN)** | **Amyloid** | C1q | 0.030 | – | n.s. |
|  |  | C3 | 0.143 | None | < .0001 |
|  |  | C3b | 0 | – | n.s. |
|  |  | Factor H | 0.008 | – | n.s. |
|  | **Tau** | C1q | 0.058 | – | n.s. |
|  |  | C3 | 0 | – | n.s. |
|  |  | C3b | 0.065 | – | n.s. |
|  |  | Factor H | 0 | – | n.s. |
| **Mild cognitive impairment (MCI)** | **Amyloid** | C1q | 0.092 | None | < .0001 |
|  |  | C3 | 0.006 | – | n.s. |
|  |  | C3b | 0.091 | None | < .0001 |
|  |  | Factor H | 0.061 | None | < .0001 |
|  | **Tau** | C1q | 0.203 | Positive | .0004 |
|  |  | C3 | 0 | – | n.s. |
|  |  | C3b | 0.174 | None | .0026 |
|  |  | Factor H | 0.062 | – | n.s. |
| **Alzheimer’s disease (AD)** | **Amyloid** | C1q | 0.146 | None | .0355 |
|  |  | C3 | 0.069 | – | n.s. |
|  |  | C3b | 0.092 | – | n.s. |
|  |  | Factor H | 0.120 | Positive | .0413 |
